# Cell-specific exon methylation and CTCF binding in neurons regulates calcium ion channel splicing and function

**DOI:** 10.1101/2019.12.15.876185

**Authors:** Eduardo Javier Lopez Soto, Diane Lipscombe

## Abstract

Cell-specific alternative splicing modulates myriad cell functions and this process is disrupted in disease. The mechanisms governing alternative splicing are known for relatively few genes and typically focus on RNA splicing factors. In sensory neurons, cell-specific alternative splicing of the presynaptic voltage-gated calcium channel *Cacna1b* gene modulates opioid sensitivity. How this splicing is regulated has remained unknown. We find that cell-specific exon DNA hypomethylation permits binding of CTCF, the master regulator of chromatin structure in mammals, which, in turn, controls splicing in noxious heat-sensing nociceptors.

Hypomethylation of an alternative exon specifically in nociceptors allows for CTCF binding, and expression of Ca_V_2.2 channels with increased opioid sensitivity. Following nerve injury, exon methylation is increased, and splicing is disrupted. Our studies define the molecular mechanisms of cell-specific alternative splicing of a functionally validated exon in normal and disease states – and reveal a potential target for the treatment of chronic pain.

**Highlights:**

- The molecular basis of cell-specific splicing of a synaptic calcium channel gene.
- Splicing controlled by cell-specific exon hypomethylation and CTCF binding.
- Peripheral nerve injury disrupts exon hypomethylation and splicing.
- Targeted demethylation of exon by dCAS9-TET modifies alternative splicing.

**GRAPHICAL ABSTRACT:** Cell-specific epigenetic modifications in a synaptic calcium ion channel gene controls cell-specific splicing in normal and neuropathic pain.
<u>In naïve animals</u>, in most neurons, *Cacna1b* e37a locus is hipermethylated (5-mC) and CTCF does not bind this locus. During splicing, e37a is skipped and *Cacna1b* mRNAs include e37b. In contrast, in *Trpv1*-lineage neurons, *Cacna1b* e37a locus is hypomethylated and is permissive for CTCF binding. CTCF promotes e37a inclusion and both *Cacna1b* e37a and e37b mRNAs are expressed. E37a confers strong sensitivity to the Ca_v_2.2 channel to inhibition by μ-opioid receptors (μOR). Morphine is more effective at inhibiting e37a-containing Ca_v_2.2 channels. <u>After peripheral nerve injury</u> that results in pathological pain, methylation level of *Cacna1b* e37a locus is increased, CTCF binding is impaired, and *Cacna1b* e37a mRNA levels are decreased. This disrupted splicing pattern is associated with reduced efficacy of morphine *in vivo*.

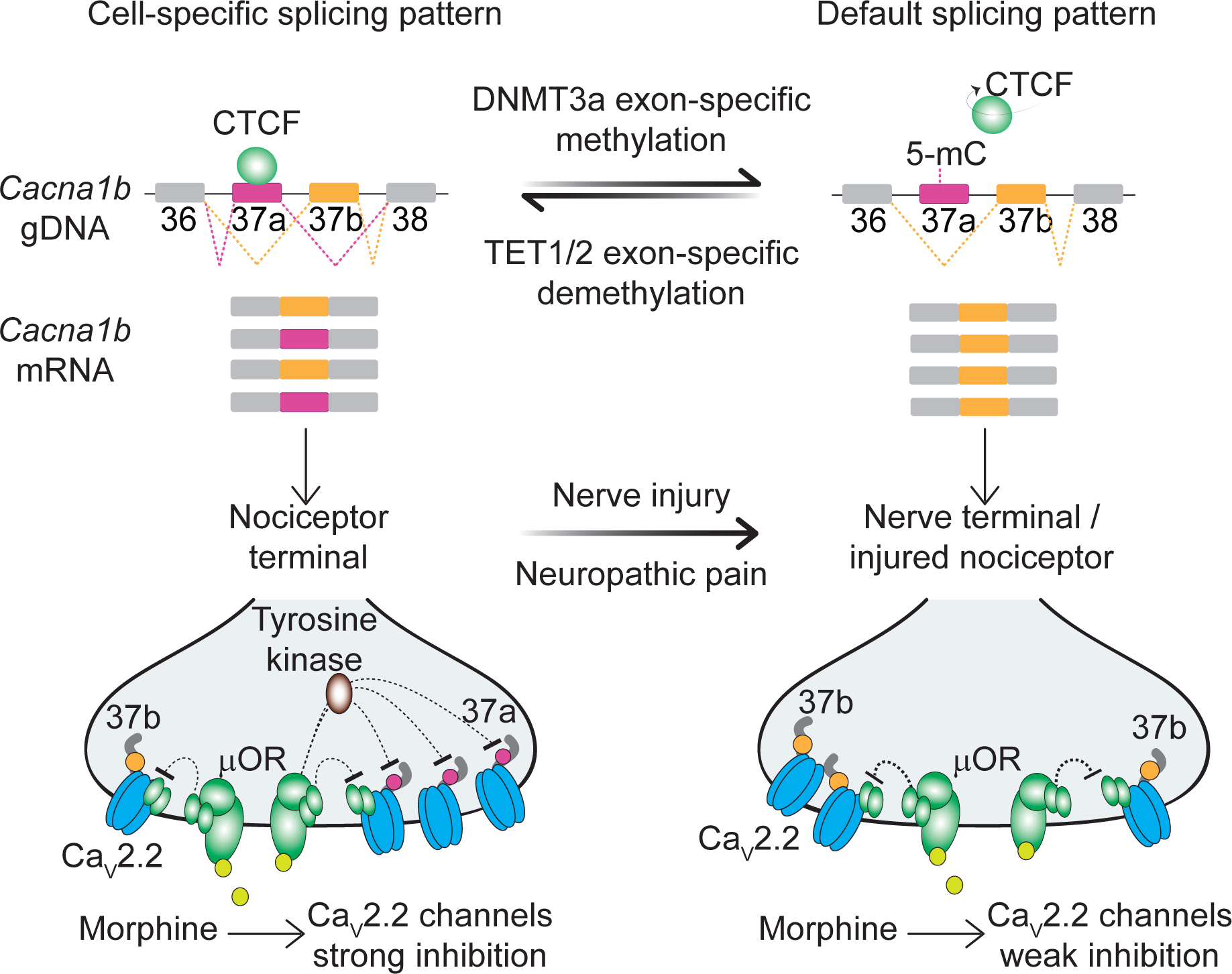

## Introduction

The precise exon composition of expressed genes defines fundamental features of neuronal function (Fiszbein and Kornblihtt, 2017; Lopez Soto et al., 2019; Ule and Blencowe, 2019). It is essential to understand the mechanisms that regulate alternative pre mRNA splicing. This dynamic process regulates exon composition for >95% of multi-exon genes according to cell-type and influenced by development, cellular activity and disease (Furlanis and Scheiffele, 2018). The cell-specific actions of RNA binding proteins are relatively well described; these splicing factors promote or repress spliceosome recruitment to pre mRNAs via *cis*-elements proximal to intron-exon splice junctions (Vuong et al., 2016). DNA binding proteins and epigenetic modifications have also been reported to alter alternative pre mRNA splicing by influencing RNA Polymerase II kinetics or splicing factor recruitment (Luco et al., 2011). However, epigenetic factors are not generally considered physiologically important regulators of cell-specific alternative pre mRNA splicing in neurons (except see (Ding et al., 2017).

Mechanisms that regulate cell-specific alternative splicing of physiologically significant events are determined for a relatively small number of exons (Furlanis and Scheiffele, 2018; Lopez Soto et al., 2019). Voltage-gated ion channel genes, essential for all electrical signaling in the nervous system, are large multi-exon genes subject to extensive alternative splicing. The cell-specific characteristics of ion channel splice isoforms determine synapse-specific release probability, neuronal firing frequencies, sensitivity to G protein coupled receptor (GPCR) modulation, subcellular targeting, and more (Gu et al., 2012; Heck et al., 2019; Lipscombe et al., 2013; Raingo et al., 2007; Thalhammer et al., 2017).

*Cacna1b* encodes the functional core of Ca_V_2.2 voltage-gated calcium channels which control presynaptic calcium entry and exocytosis at mammalian synapses. Numerous drugs and neurotransmitters downregulate synaptic transmission via GPCR that act on Ca_V_2.2 channels (Huang and Zamponi, 2017). *Cacna1b* generates Ca_V_2.2 splice isoforms with unique characteristics, including sensitivity to GPCRs, that underlie their functional differences across the nervous system (Allen et al., 2010; Bunda et al., 2019; Gandini et al., 2019; Macabuag and Dolphin, 2015; Marangoudakis et al., 2012; Raingo et al., 2007). The best characterized of these involves a mutually exclusive exon pair (e37a and e37b). Ca_V_2.2 channels that contain e37a, in place of the more prevalent e37b, are expressed in a subset of nociceptors and they are especially sensitive to inhibition by μ-opioid receptors (Bell et al., 2004; Castiglioni et al., 2006; Macabuag and Dolphin, 2015; Raingo et al., 2007). Cell-specific inclusion of e37a enhances morphine analgesia *in vivo*, and disruption of this splicing event following nerve injury reduces the action of morphine (Altier et al., 2007; Andrade et al., 2010; Jiang et al., 2013).

Here we report the molecular mechanisms that regulate this physiologically significant splicing event, and the surprising finding that the ubiquitous 11-zinc finger CCCTC binding factor (CTCF), the master regulator of chromatin architecture in mammals, and CpG methylation are critical for cell-specific expression of *Cacna1b* e37a in noxious heat sensing nociceptors. Our studies define, for the first time, the mechanisms of cell-specific alternative splicing of a synaptic ion channel gene exon in normal and in disease states.

## RESULTS

### The ubiquitous DNA binding protein CTCF binds the *Cacna1b* e37a locus

To screen for factors governing cell-specific exon selection at *Cacna1b* e37 loci we searched publicly available databases for RNA and DNA binding protein associated with this region (Fig. 1A). We found no evidence for any RNA binding protein associating with *Cacna1b* e37a or e37b, based on analyses of cross-linking immunoprecipitation following by sequencing (CLIP-seq) data. However, we observed a robust chromatin immunoprecipitation followed by sequencing (ChIP-seq) signal for the zinc finger DNA binding protein CCCTC-binding factor (CTCF) that overlaps the *CACNA1B* e37a locus in ~ 50% of human cell lines (27 of 50; 9 of 50 tracks are shown in Fig. 1B) (Consortium, 2012). None of the 50 tracks contained a ChIP-seq CTCF signal associated with *CACNA1B* e37b (Fig. 1B).

**Figure 1.**
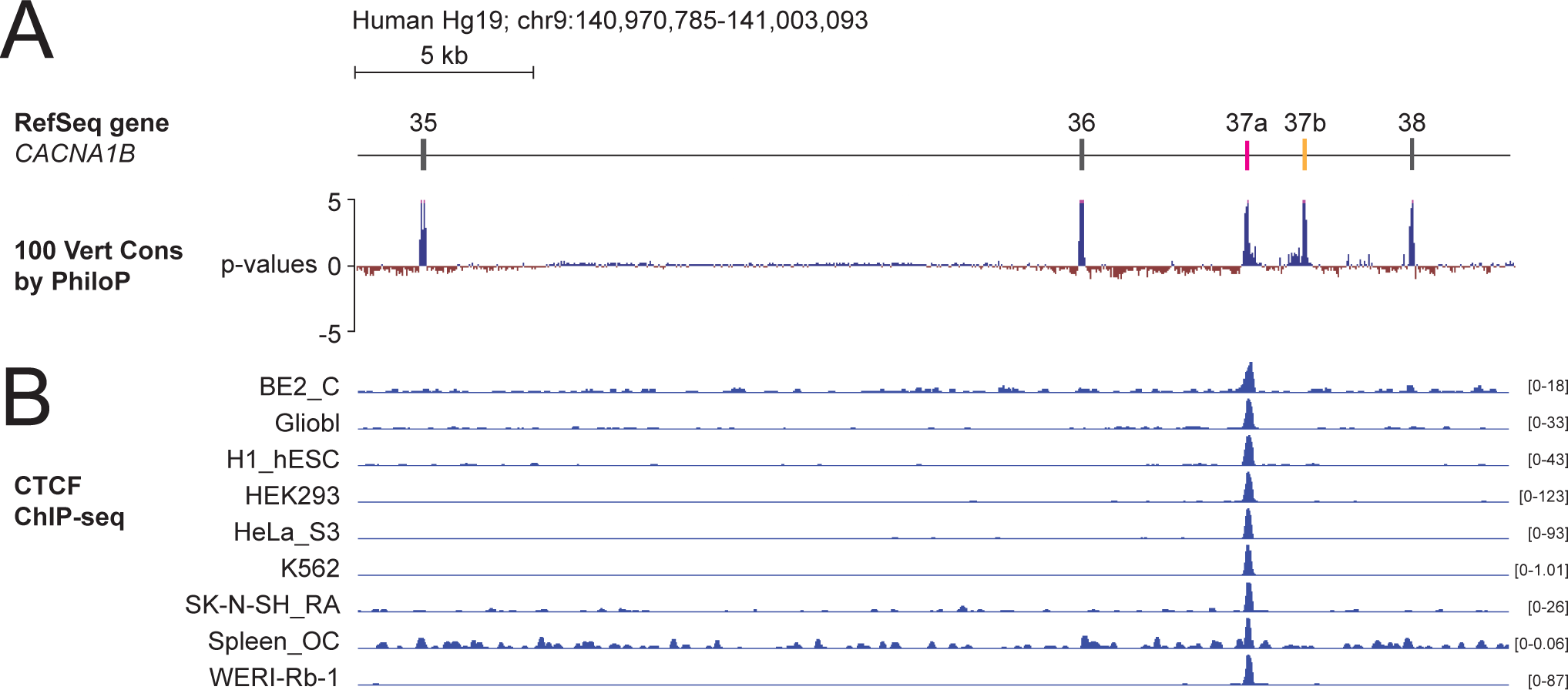
The DNA binding protein CTCF binds *Cacna1b* e37a but not e37b *in vivo*. (A) The 100 vertebrate basewise conservation track for ~ 30 kb region of *Cacna1b* (Hg19; chr9:104,970,785-141,003,093). Five conserved elements align to e35, e36, e37a, e37b, and e38. (B) ChIP-seq signals for CTCF binding in 9 different human cell lines are aligned to *CACNA1B* region in *A*. Y-axes for ChIP-seq tracks are scaled to the maximum signal within the selected region. In total, there is CTCF signal at *CACNA1B* e37a in 27 of 50 human cell lines. Link to the UCSC genome output (https://genome.ucsc.edu/s/ejlopezsoto/Cacna1b%20e35%20to%20e38%20conservation%20track) (Consortium, 2012).

In addition to CTCF, four other DNA binding proteins associate with *CACNA1B* e37a but in far fewer cell lines compared to CTCF (Supplemental Fig. 1). Of these, RAD21 (3 of 27 cell lines) and SMC3 (1 of 27 cell lines) are often found in a complex with CTCF (Zhang et al., 2018); CTCFL (1 of 27 cell lines) is a CTCF-like testes-specific DNA binding protein (Loukinov et al., 2002), and CEBPB (3 of 27 cell lines) is associated with gene enhancers (Supplemental Fig. 1A) (Nerlov, 2007). We focused on CTCF as the most likely factor involved in enhancing *CACNA1B* e37a inclusion during pre-mRNA splicing given these data, and because CTCF has been proposed to influence exon recognition in *CD45* (Shukla et al., 2011).

CTCF is ubiquitously expressed in the bilaterian phyla (Heger et al., 2012) and widely recognized as the master organizer of chromatin in mammals (Ong and Corces, 2014). Notably, CTCF was proposed as a regulator of alternative splicing in immune cells (Ruiz-Velasco et al., 2017; Shukla et al., 2011), although a role for CTCF in regulating cell-specific splicing has not been proposed in neurons.

Several observations suggested to us that CTCF might be the key factor promoting *Cacna1b* e37a recognition in neurons: CTCF binding was robust in many, but not all human cell lines (Fig. 1B); *Cacna1b* e37a contains a highly conserved consensus CTCF binding motif that is not present in e37b (Fig. 2A); and it associates with mouse *Cacna1b* e37a but not e37b, which share 60% nucleotide identity (Fig. 2A, 2B). We therefore set out to test this hypothesis *in vitro* and *in vivo*.

**Figure 2.**
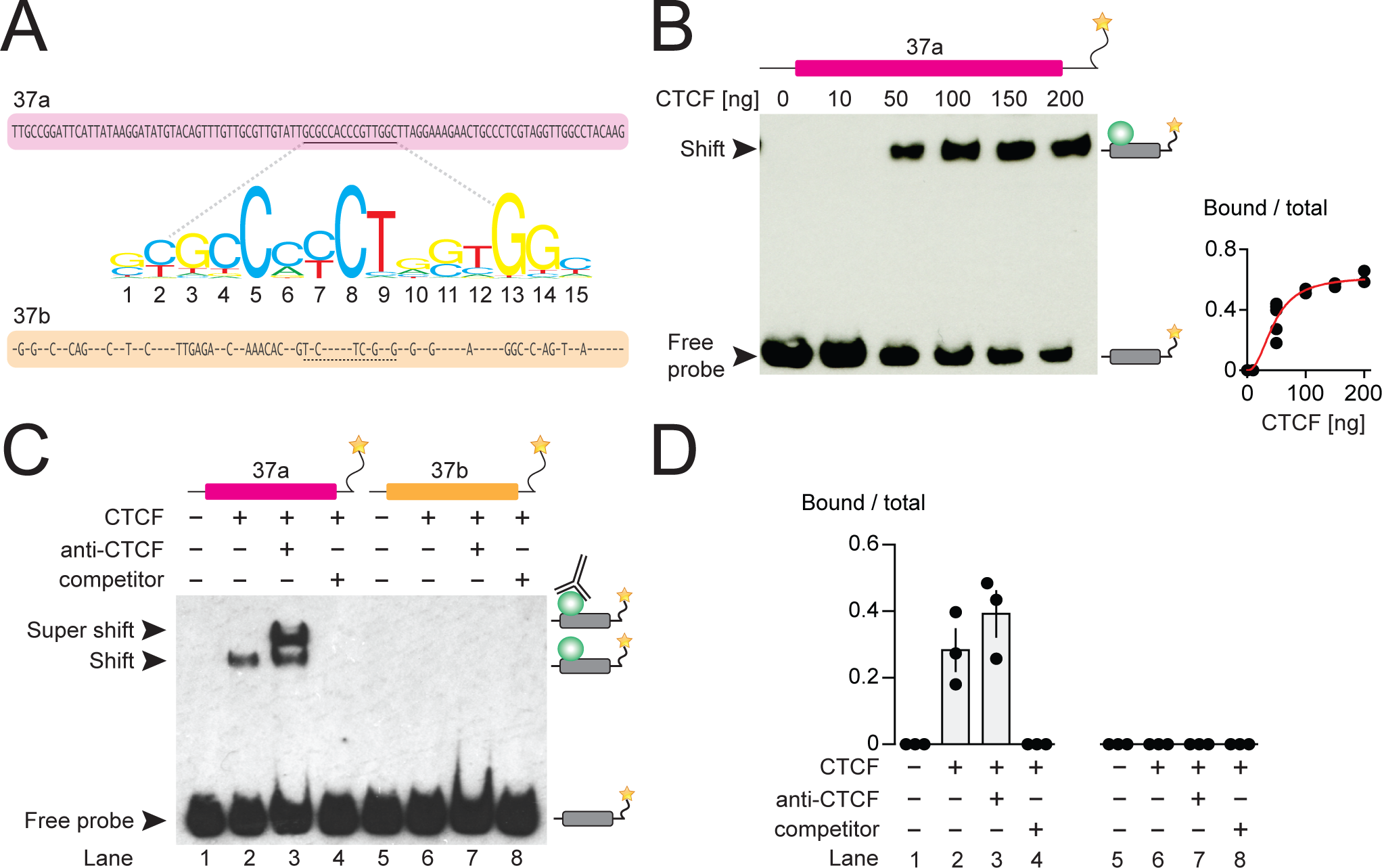
CTCF binds mouse *Cacna1b* e37a *in vitro*. (A) Mouse *Cacna1b* e37a and e37b sequences are ~60% identical. Nucleotides conserved across exons are shown as dashes in the e37b sequence. A 15 nucleotide CTCF binding motif in e37a (underlined) is not present in e37b. (B) Gel shift assay for CTCF and a *Cacna1b* e37a DNA probe (139 bp) that also contains 5’ and 3’ flanking intron sequences alone (0 ng) or with increasing concentrations of recombinant CTCF (10-200 ng). Quantification of CTCF bound to e37a probe relative to total e37a probe at different CTCF concentrations (red line represents fitted Hill equations r2 = 0.9338; *Kd* = 0.0461 ± 0.0055 ng) (n = at least 2 per CTCF concentration). (C) Gel shift assay for CTCF and DNA probes (139 bp) containing either *Cacna1b* e37a or e37b alone (lanes 1 and 5) or preincubated with 50 ng recombinant CTCF (lanes 2 and 6), CTCF plus 0.5 μg anti-CTCF (lanes 3 and 7), or CTCF plus 1000-fold excess unlabeled e37a or e37b probes (lanes 4 and 8). (D) Quantification of CTCF bound to e37 probes relative to total e37a or e37b probe (ANOVA *P* value = 0.0006 for e37a probe, Dunnett’s multicomparison *P* values for lane 1 vs 2 = 0.0067, lane 1 vs 3 = 0.0009, lane 1 vs 4 = 0.9999) (n = 3 per condition). Biological replicates represent independent gel shift assays.

### CTCF binds *Cacna1b* e37a *in vitro*

We investigated direct and specific binding of CTCF to *Cacna1b* e37a *in vitro* using the electrophoretic mobility shift assay. Recombinant CTCF bound to an e37a-containing DNA probe in a concentration manner (Fig. 2B) but CTCF failed to bind an e37a-containing RNA probe (Supplemental Fig. 2A). To establish specificity of binding, we incubated labelled e37a and e37b DNA probes (139 bp) alone; with recombinant CTCF; with CTCF plus CTCF monoclonal antibody; or with CTCF plus unlabeled probe (Fig. 2C, 2D). In the presence of 50 ng recombinant CTCF, ~30-40% of the e37a DNA probe migration was slowed, relative to no CTCF, as indicated by the presence of a second band which was shifted to higher molecular weights (Fig. 2C, 2D; compare lanes 1 and 2). Based on the following observations we conclude that CTCF binds with high specificity to *Cacna1b* e37a but not *Cacna1b* e37b: 1) CTCF bound e37a but not e37b DNA probes when studied using the same conditions (Fig. 2C, 2D; compare lanes 2 and 6); 2) the appearance of a third super-shifted band when e37a probe is incubated with both recombinant CTCF and a CTCF monoclonal antibody (Fig. 2C, 2D; lane 3); 3) total displacement of CTCF binding from the labeled e37a DNA probe on addition of unlabeled e37a DNA probe (Fig. 2C, 2D; lane 4); and CTCF binding to e37a DNA is concentration dependent (Fig. 2B). We also tested the ability of various truncated e37a DNA probes to bind CTCF, but CTCF binding was reduced in all truncated constructs (Supplemental Fig. 2B). This is consistent with other reports that CTCF binding to DNA is strongly influenced by several factors including probe length (Fitzpatrick et al., 2007).

### CTCF binds *Cacna1b* e37a locus *in vivo* and modifies e37a splicing in neuronal cells

Having shown that CTCF binds directly and specifically to *Cacna1b* e37a locus *in vitro*, but not e37b, we next tested if CTCF levels influence *Cacna1b* e37a splicing in neuronal cells. We used the rat DRG/mouse neuroblastoma hybrid F11 cell line which expresses *Cacna1b* (Allen et al., 2017), to determine if CTCF binds *Cacna1b* e37a locus in a cellular context, and to test if CTCF levels influence *Cacna1b* e37a splicing. CTCF is localized to cell nuclei in F11 cells (Fig. 3A). We found that CTCF binds *Cacna1b* e37a locus in F11 cells by CTCF ChIP-qPCR (Fig. 3B). By contrast, CTCF does not bind e37b locus (Fig. 3B). Our findings in F11 cells, that CTCF associates with e37a but not e37b, are consistent with ChIP-seq data analyzed in 27 different human cell lines (see Fig. 1).

**Figure 3.**
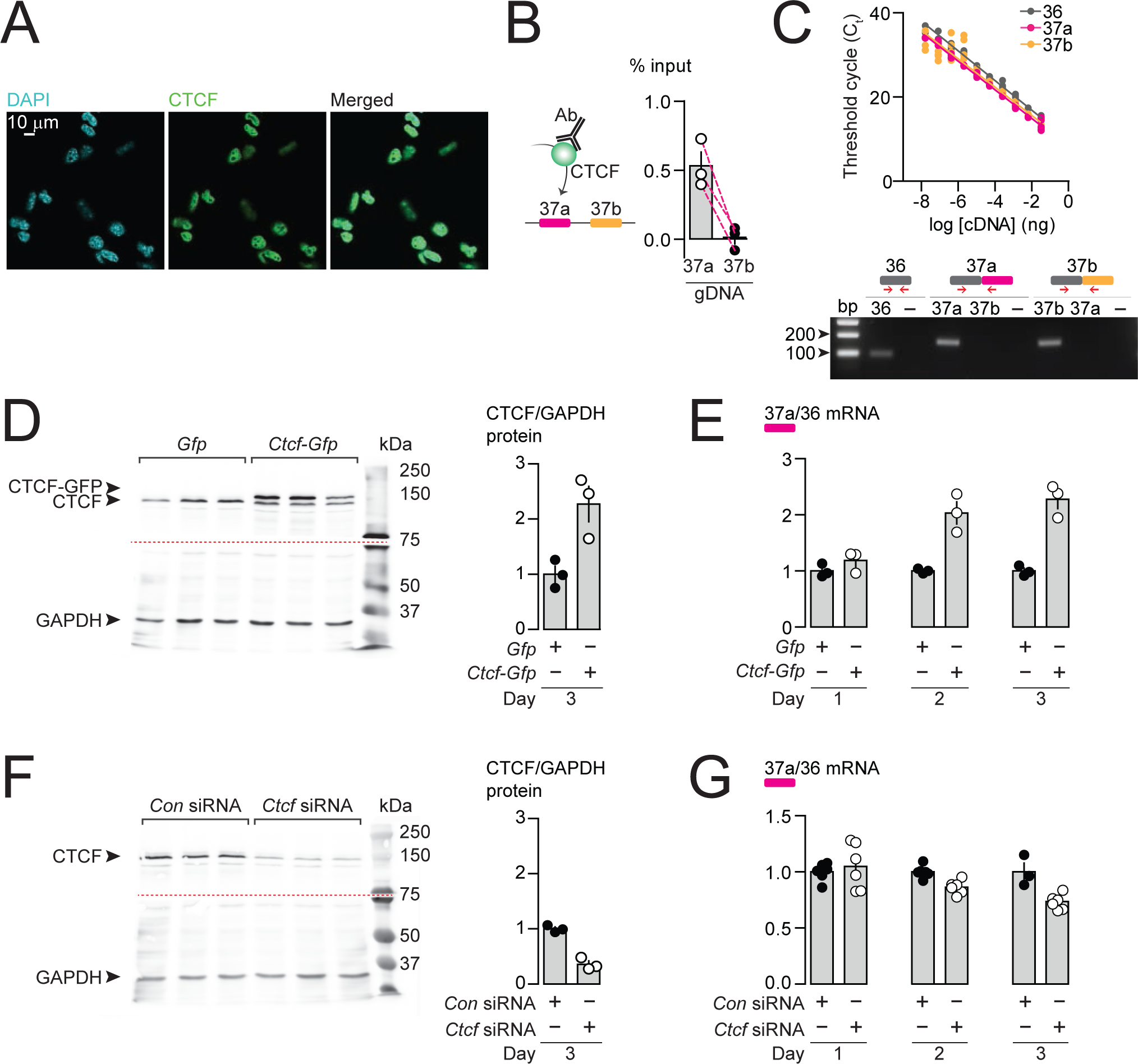
CTCF binds *Cacna1b* e37a locus in DRG-derived F11 cells and influences e37a splicing. (A) CTCF is expressed in F11 cells and is localized to cell nuclei. Fluorescence images show DAPI (nuclear maker), anti-CTCF, and overlay. (B) CTCF ChIP-qPCR targeted to *Cacna1b* e37 loci in F11 cells (Paired t-test, *P* value = 0.0274 for e37a vs e37b) (n = 3 per condition). (C) Efficiency and specificity of qPCR primers. *Cacna1b* e36, e37a and e37b-specific primers amplify DNA with similar log-linear relationships between 1 × 10^−8^ and 1.5 × 10^−1^ ng cDNA (*upper panel*) (gray, pink and orange lineal regression lines for e36, e37a and e37b data sets have similar slopes; Fisher’s tests *P* value = 0.4383). qPCR primer specificity was confirmed by using *Cacna1b* e37a and e37b cDNA clones (*lower panel*). (D) Western blot from F11 cells expressing control *Gfp* or *Ctcf-Gfp* cDNA 3 days after transfection. In control cells, anti-CTCF identifies a single band at ~135 kDa. Whereas, in cells overexpressing *Ctcf-Gfp*, anti-CTCF identifies two bands, endogenous ~135 kDa and a second ~160 kDa band consistent with the expected size of the CTCF-GFP fusion protein. Transfer membrane was cut at ~ 75 kDa (dotted red line) and the lower part treated with anti-GAPDH to measure GAPDH levels for protein expression and loading reference. CTCF protein levels relative to GAPDH (t-test *P* value = 0.0227 for *Gfp* vs *Ctcf-Gfp*) (n = 3 per condition). (E) qRT-PCR of *Cacna1b* e37a relative to e36 in F11 cells overexpressing *Gfp* or *Ctcf -Gfp* cDNA over three days (t-test *P* values = 0.2263 for *Gfp* vs *Ctcf -Gfp* for day 1; *P* = 0.0065 for day 2; and *P* = 0.0019 for day 3) (n = 3 per condition). (F) Western blot from F11 cells expressing control siRNA (*Con*) or *Ctcf* siRNA 3 days after transfection. In control cells, anti-CTCF identifies endogenous CTCF at ~135 kDa, and reduced CTCF levels in cells expressing *Ctcf* siRNA. Transfer membrane was cut at ~ 75 kDa (dotted red line) and the lower part treated with anti-GAPDH to measure GAPDH levels for protein expression and loading reference. CTCF protein levels relative to GAPDH (t-test *P* value = 0.0008 for *Con* vs *Ctcf* siRNA) (n = 3 per condition). (G) qRT-PCR of *Cacna1b* e37a relative to e36 in F11 cells expressing control siRNA (*Con*) or *Ctcf* siRNA over three days (t-test *P* values = 0.6035 for *Con* vs *Ctcf* siRNA for day 1; *P* = 0.0022 for day 2; and *P* = 0.0046 for day 3) (n = at least 3 per condition). Biological replicates represent independent cell cultures, treatment and transfections.

We next tested if CTCF levels in F11 cells influence the expression of *Cacna1b* e37a mRNAs. We conducted high-efficient qPCR using primer pairs for e37a, e37b, and e36 (constitutive exon), which were optimized and matched for specificity and accuracy, to quantify mRNAs (Fig. 3C). We either overexpressed CTCF, or knocked down CTCF in F11 cells (Fig. 3D, 3F) and then quantified levels of *Cacna1b* e37a mRNAs, relative to total *Cacna1b* mRNAs (e36; Fig. 3E, 3G). When overexpressed (Fig. 3D, 3E), recombinant CTCF tagged with GFP induced a ~2-fold increase in *Cacna1b* e37a levels within 2 days of cDNA transfection, as compared to cells expressing *Gfp* alone (Fig. 3E). siRNA targeting *Ctcf* in F11 cells (Fig. 3F, 3G) had the expected opposite effect; levels of e37a-containing *Cacna1b* mRNAs were ~30% reduced 3 days after transfecting *Ctcf* siRNA relative to control siRNA (Fig. 3G). Our findings show that CTCF promotes e37a inclusion in *Cacna1b* mRNAs, and that CTCF levels are rate limiting for *Cacna1b* e37a splicing in neuronal cells.

### 5-mC DNA modification controls CTCF binding and *Cacna1b* e37a inclusion

CTCF binding to gDNA is reported to be inhibited by 5-methylcytosine (5-mC) within CTCF binding motif and interferes on control of alternative splicing in *CD45* (Hashimoto et al., 2017; Marina et al., 2016; Shukla et al., 2011). We used pharmacological agents to alter global gDNA 5-mC, assessed their impact on *Cacna1b* e37a splicing and e37a locus CpG 5-mC in F11 cells (Fig. 4; Supplemental Fig. 3).

**Figure 4.**
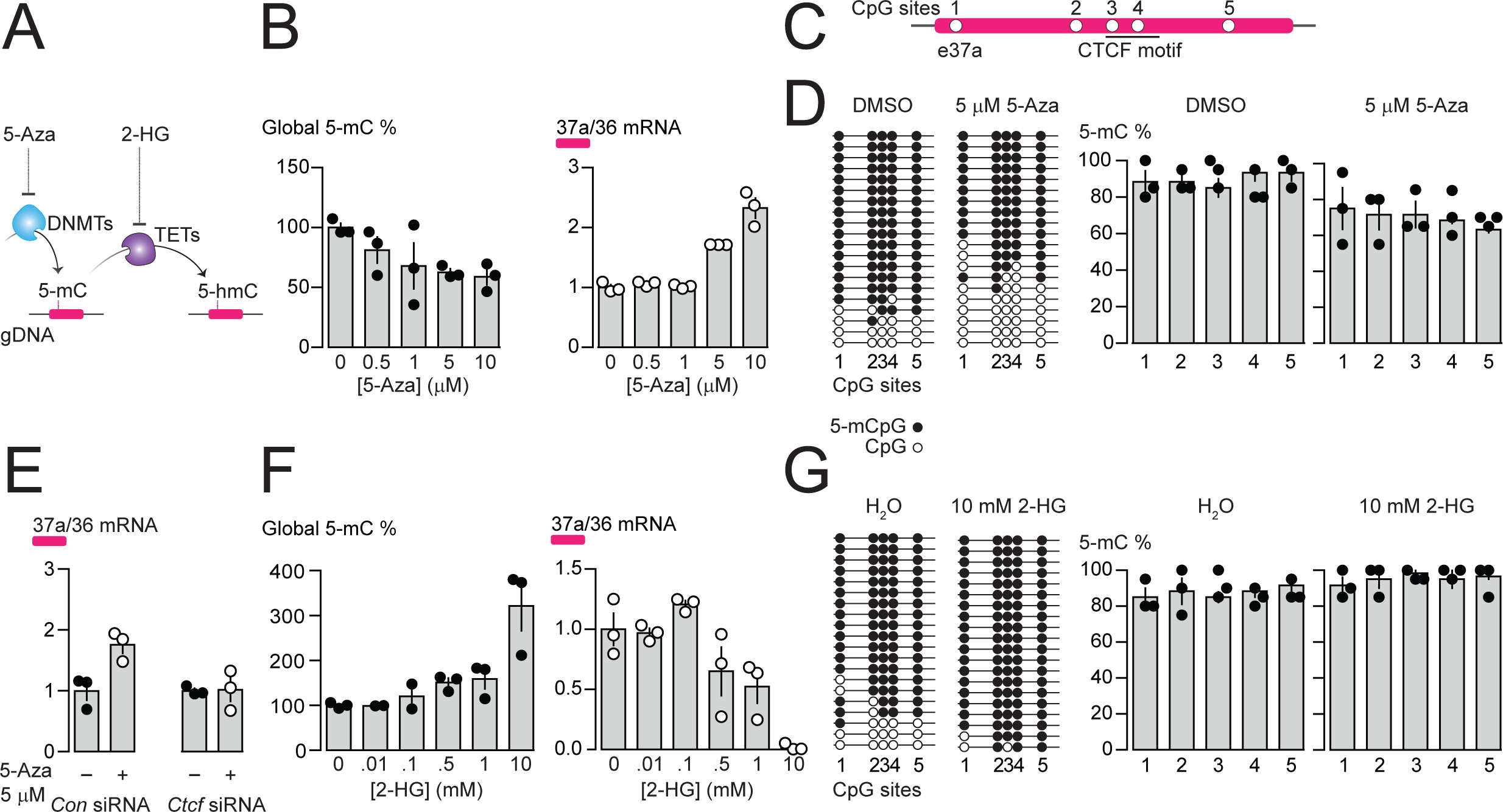
Pharmacological manipulation of gDNA methylation in F11 cells alters 5-mC levels at CpG sites in *Cacna1b* e37a locus and e37a inclusion. (A) 5-Azacytidine (5-Aza) inhibits the activity of DNA methyltransferases (DNMTs), and 2-hydroxyglutarate (2-HG) inhibits the activity of ten-eleven translocation proteins (TETs). (B) Global 5-mC levels in F11 cells after 3 day exposure to different concentrations of 5-Aza (*left*) (ANOVA *P* value = 0.056, Dunnett’s multicomparison *P* values for 0 vs 0.5 μM = 0.5371, 0 vs 1 μM = 0.1333, 0 vs 5 μM = 0.0688, 0 vs 10 μM = 0.0421, and 0 vs 10 μM = 0.0280) and associated qRT-PCR of *Cacna1b* e37a relative to e36 levels (*right*) (ANOVA *P* value = <0.0001, Dunnett’s multicomparison *P* values for 0 vs 0.5 μM = 0.9874, 0 vs 1 μM = 0.9998, 0 vs 5 μM = 0.0015, 0 vs 10 μM = <0.0001) (n= 3 per condition). (C) Location of 5 CpG sites in *Cacna1b* e37a locus (open circles) and CTCF binding motif (underlined). (D) 5-mC levels at CpG sites in *Cacna1b* e37a locus in control F11 cells (DMSO, *left*) and after 3-day treatment with 5 μM 5-Aza (*right*). Methylated (5-mCpG, filled circle) and unmethylated (CpG, open circle) sites are shown for independent clones ordered from most to least methylated. 5-mC levels for both conditions (two-way ANOVA *P* value = 0.0260 for treatment factor) (n = 3 per condition). (E) qRT-PCR of *Cacna1b* e37a relative to e36 in F11 cells in the absence and after 3-day treatment with 5 μM 5-Aza for cells expressing control siRNA (*Con*) or *Ctcf* siRNA (t-test *P* value = 0.0219 for untreated vs 5-Aza with *Con* siRNA; and = 0.9303 for untreated vs 5-Aza with *Ctcf* siRNA) (n = 3 per condition). (F) Global 5-mC levels in F11 cells following 1-day exposure to different concentrations of 2-HG (*left*) (ANOVA *P* value = 0.0022, Dunnett’s multicomparison *P* values for 0 vs 0.01 mM = >0.9999, 0 vs 0.1 mM = 0.9878, 0 vs 0.5 mM = 0.6062, 0 vs 1 mM = 0.4755, and 0 vs 10 mM = 0.0010), and associated qRT-PCR of *Cacna1b* e37a relative to e36 levels (*right*) (ANOVA *P* value = 0.0001, Dunnett’s multicomparison *P* values for 0 vs 0.01 mM = 0.9997, 0 vs 0.1 mM = 0.6026, 0 vs 0.5 mM = 0.1799, 0 vs 1 mM = 0.0487, and 0 vs 10 mM = 0.0003) (n= 3 per condition). (G) 5-mC levels at CpG sites in *Cacna1b* e37a locus in control F11 cells (DMSO, *left*) and after 1-day treatment with 10 mM 2-HG (*right*). Methylated (5-mCpG, filled circle) and unmethylated (CpG, open circle) sites are shown for independent clones ordered from most to least methylated. 5-mC levels for both conditions (two-way ANOVA *P* value = 0.0745 for treatment factor) (n = 3 per condition). Biological replicates represent independent cell cultures, treatment and transfections.

We used the pharmacological inhibitor of DNA methyltransferases (DNMTs), 5-Azacytidine (5-Aza), to reduce overall 5-mC levels in F11 cells (Fig. 4A). Increasing concentrations of 5-Aza decreased global gDNA 5-mC levels in F11 cells to ~60 % of control (Fig. 4B), without altering CTCF protein levels or global levels of 5-hydroxymethylcytosine (5-hmC) (Supplemental Fig. 3B, 3C). Reduced 5-mC levels induced by 5-Aza were associated with a concentration dependent increase in *Cacna1b* e37a mRNA levels in F11 (Fig. 4B). We used 5 μM 5-Aza for 3 days in subsequent experiments to maximize effects on 5-mC levels, while limiting cell toxicity (Fig. 4D, 4E); higher concentrations of 5-Aza were cytotoxic (Supplemental Fig. 3A).

By targeted bisulfite sequencing, we found that 5 μM 5-Aza treatment in F11 cells, which induced a >2-fold increase in *Cacna1b* e37a mRNA levels, reduced 5-mC levels of the 5 CpG sites in *Cacna1b* e37a locus by ~10% relative to control (DMSO) (Fig. 4C, 4D). We also found that the ability of 5-Aza to promote *Cacna1b* e37a inclusion in F11 cells depended on CTCF availability. In cells expressing *Ctcf* siRNA, 5-Aza had no effect on e37a inclusion (Fig. 4E). These experiments reinforce our hypothesis that CTCF is a factor in *Cacna1b* e37a recognition and that it promotes e37a inclusion during *Cacna1b* pre-mRNA splicing.

Next, we used the global pharmacological inhibitor of ten eleven translocase enzymes (TETs), 2-hydroxyglutarate (2-HG), to increase overall 5-mC levels in F11 cells (Fig. 4A). Increasing concentrations of 2-HG increased global DNA 5-mC levels in F11 cells (Fig. 4F), reduced levels of 5-hydroxymethylcytosine (5-hmC), but did not alter CTCF protein levels (Supplemental Fig. 3D, 3E). 10 mM 2-HG treatment for 1 day induced ~ 3-fold increase in global 5-mC methylation in F11 cells (Fig. 4F) and, under these conditions, reduced *Cacna1b* e37a mRNAs to almost undetectable levels (Fig. 4F). By targeted bisulfite sequencing, we found that 10 mM 2-HG treatment in F11 cells induces a slight increase of ~ 10% in 5-mC levels at the 5 CpG sites in *Cacna1b* e37a locus relative to control (Fig. 4G).

In the above experiments, global pharmacological manipulation of 5-mC gDNA reveals a negative correlation between 5-mC at the *Cacna1b* e37a locus and *Cacna1b* e37a mRNA levels in F11 cells. Our findings support the hypothesis that 5-mC within *Cacna1b* e37a locus occludes CTCF binding and impairs CTCF-mediated exon recognition during alternative pre-mRNA splicing.

### DNMT3a, TET1 and TET2 control CTCF actions on *Cacna1b* e37a splicing

To identify the specific enzymes that regulate the 5-mC within *Cacna1b* e37a locus in F11 cells, we knocked down individual DNMTs, and overexpressed cDNAs encoding known TETs, implicated in dynamic regulation of gDNA methylation in neurons (Chen et al., 1991; He et al., 2011) and then quantified the impact on *Cacna1b* e37a expression by qRT-PCR (Fig. 5A, 5B). We found that *Dnmt3a* siRNA (3a), but not *Dnmt1* (1) or *Dnmt3b* siRNAs (3b), promoted >2-fold increase *Cacna1b* e37a inclusion, compared to control siRNA in F11 cells (Fig. 5A); and that overexpressed cDNAs encoding *Tet1* (1) and *Tet2* (2), but not *Tet3* (3) cDNAs, promoted ~2-fold increase *Cacna1b* e37a inclusion in mRNA in F11 cells compared to control (Fig. 5B).

**Figure 5.**
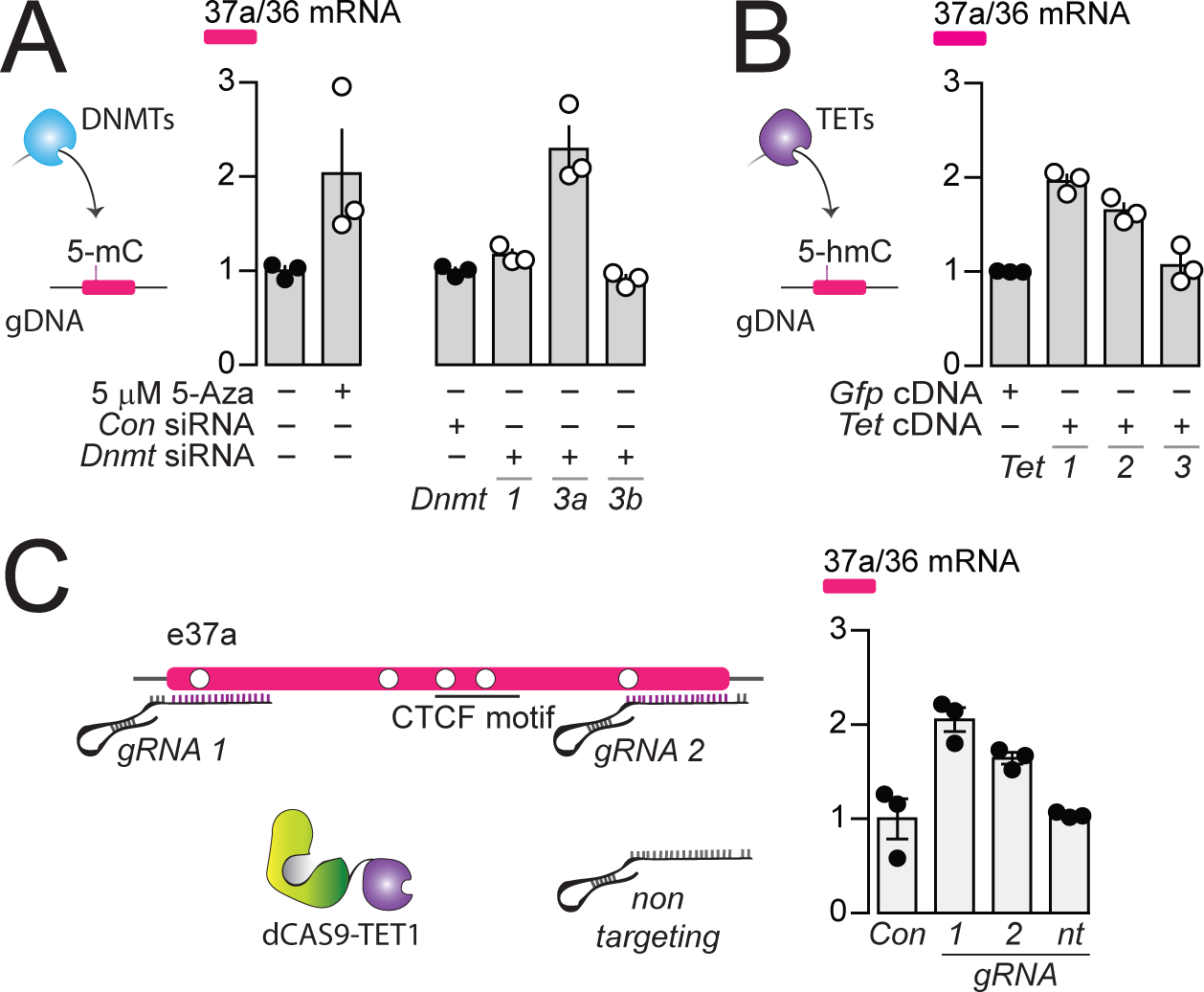
The DNA methyltransferase 3a inhibits, and ten-eleven translocating protein 1 and 2 promote *Cacna1b* e37a inclusion. (A) DNA methyltransferases (DNMTs) methylate CpG sites in gDNA. qRT-PCR of *Cacna1b* e37a relative to e36 levels in F11 cells in the absence and after 3-day treatment with 5 μM 5-Aza (t-test *P* value = 0.0923), and in cells expressing control siRNAs (*Con*) or siRNAs targeting *Dnmt1*, *Dnmt3a*, or *Dnmt3b* after 3 days (ANOVA *P* value = 0.0002, Dunnett’s multicomparison *P* values for *Con* vs *Dnmt1* siRNA = 0.6925, *Con* vs *Dnmt3a* siRNA = 0.0003, and *Con* vs *Dnmt3b* siRNA = 0.9226) (n = 3 per condition). (B) Ten-eleven translocating proteins (TETs) promote demethylation of CpG sites in gDNA. qRT-PCR of *Cacna1b* e37a relative to e36 levels in F11 cells expressing control *Gfp*, *Tet1*, *Tet2*, or *Tet3* cDNAs for 3 days (ANOVA *P* value = <0.0001, Dunnett’s multicomparison *P* values for *Gfp* vs *Tet1* cDNA = <0.0001, *Gfp* vs *Tet2* cDNA = 0.0008, and *Gfp* vs *Tet3* cDNA = 0.8676) (n = 3 per condition). (C) dCAS-targeting strategy to localize TET1 to *Cacna1b* e37a locus using guide RNAs (gRNA). qRT-PCR of *Cacna1b* e37a relative to e36 levels in F11 cells expressing dCas9-Tet1 alone (*Con*) or with gRNAs to the 5’ (*1*) or 3’ (*2*) ends of *Cacna1b* e37a locus or non-targeting gRNA (*nt*) (ANOVA *P* value = 0.0010, Dunnett’s multicomparison *P* values for *Con* vs *gRNA 1* = 0.0010, *Con* vs *gRNA 2* = 0.0186, and *Con* vs *gRNA nt* = 0.9914) (n = 3 per condition). Biological replicates represent independent cell cultures, treatment and transfections.

We next employed deactivated CAS9 (dCAS9) to target TET1 directly to *Cacna1b* e37a locus. We target-optimized guide RNA (gRNA) sequences to 5’ and 3’ regions of *Cacna1b* e37a locus to localize dCAS9-TET1 and measured *Cacna1b* e37a inclusion by qRT-PCR in F11 cells (Fig. 5C). Both gRNA-dCAS9-TET1 complexes increased *Cacna1b* e37a expression in F11 by ~2-fold compared to F11 cells expressing *dCas9-Tet1* alone (Con) or to F11 cells expressing a non-targeting gRNA (nt) (Fig. 5C).

Our results support a model in which *Cacna1b* e37a inclusion during alternative splicing is inhibited by DNMT3a and promoted by TET1 and TET2.

### *Cacna1b* e37a is expressed in *Trpv1*-lineage sensory neurons of mouse dorsal root ganglia

In previous studies, we showed that *Cacna1b* e37a mRNAs are enriched in DRG of rat and, by single cell RT-PCR coupled to electrophysiological analyses, we found further enrichment in a subset of capsaicin-responsive nociceptors (Bell et al., 2004). Here we extend these studies and use a mouse strain expressing the *TdTomato* or *Yfp* reporter in *Trpv1*-lineage neurons for fluorescence-activated cell sorting (FACS). By endpoint RT-PCR, we found that *Cacna1b* e37a mRNAs are exclusive to *Trpv1*-lineage neurons of DRG (Fig. 6A), whereas *Cacna1b* e37b mRNAs are expressed in both *Trpv1*-lineage and non *Trpv1*-lineage cells (Fig. 6A).

**Figure 6.**
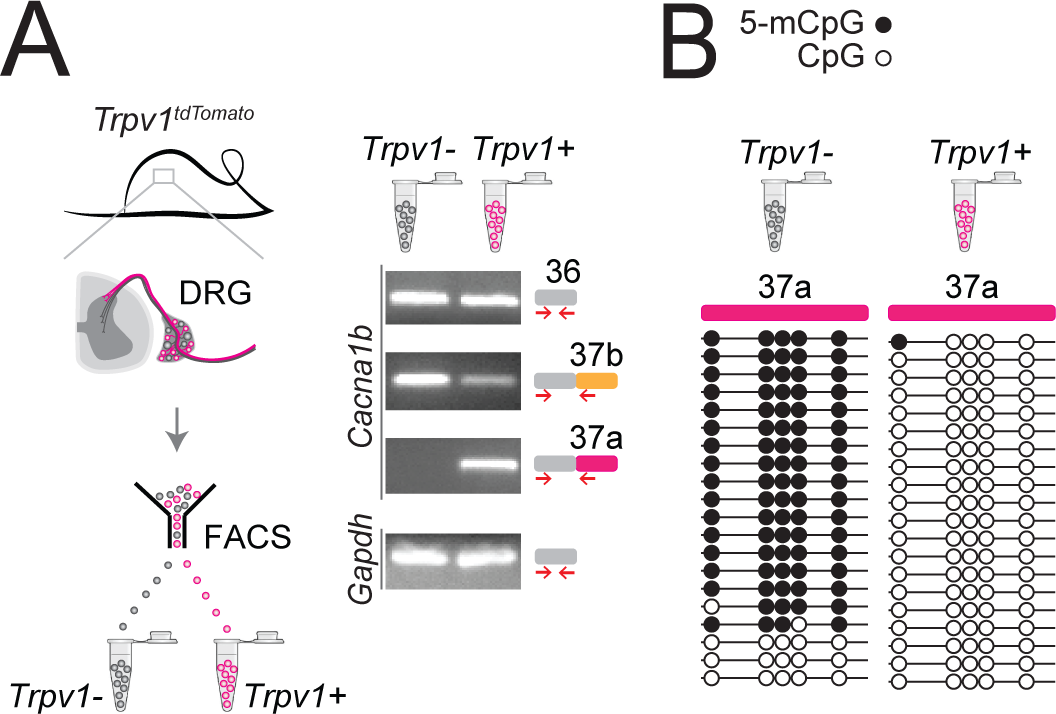
Cell-specific *Cacna1b* e37a inclusion and methylation in *Trpv1*-lineage nociceptors. (A) *Trpv1*^*Cre*^ mouse strain crossed to *lox-STOP-lox*^*TdTomato*^ expressed the *TdTomato* reporter in *Trpv1*-lineage neurons (*Trpv1*^*TdTomato*^). DRG cells were separated into *Trpv1*-lineage and non-*Trpv1*-lineage cells by fluorescence-activated cell sorting (FACS). Using specific primer pairs RT-PCR products amplified: constitutive *Cacna1b* e36, alternative spliced e37a and e37b, and *Gapdh* in *Trpv1*-lineage and non *Trpv1*-lineage cells for all primer pairs except for non *Trpv1*-lineage cells that failed to amplify from e37a-specific primers. Data shown are representative of a biological replicate from at least 3 different FACS-separated DRGs from 3 animals. (B) Methylation state of 5 CpG sites in *Cacna1b* e37a locus in non *Trpv1*-lineage (left) and in *Trpv1*-lineage neurons (*right*). Methylated (5-mCpG, filled circle) and unmethylated (CpG, open circle) sites are shown for independent clones ordered from most to least methylated. Each set of sequences represents data pooled from at least 10 FACS-separated DRG cells from 1 animal. These results are confirmed in Fig. 7D for 6 additional animals.

The striking cell-specific expression pattern in *Trpv1*-lineage, but not in non *Trpv1*-lineage neurons, offered a way to test our hypothesis *in vivo*: If 5-mC CpG of *Cacna1b* e37a locus is important for e37a recognition, then *Trpv1*-lineage neurons should have reduced 5-mC levels relative to non *Trpv1*-lineage DRG cells in *Cacna1b* e37a locus. We used targeted bisulfite sequencing and found that all 5 CpG sites in *Cacna1b* e37a locus in non *Trpv1*-lineage cells are methylated in the majority of sequences (Fig 6B). By contrast, of 20 independent clones derived from *Trpv1*-lineage neurons, only one CpG site in *Cacna1b* e37a locus, in one sequence contained 5-mC (Fig 6B).

Our results show that 5-mC CpGs of *Cacna1b* e37a locus are cell-specific – CpG sites in e37a are hypomethylated in *Trpv1*-lineage neurons relative to the close to fully methylated state of e37a locus in non *Trpv1*-lineage cells. These data provide strong independent support for our hypothesis, and they suggest that cell-specific control of local methylation regulates the level of *Cacna1b* e37a inclusion during pre-mRNA splicing.

### Peripheral nerve injury increases 5-mC within *Cacna1b* e37a locus in Trpv1-lineage neurons

We showed previously that *Cacna1b* e37a mRNA levels decrease in rat DRG following peripheral nerve injury, and this disruption in splicing is linked to reduced efficacy of morphine analgesia (Altier et al., 2007; Jiang et al., 2013). We therefore tested if nerve injury-induced disruption of *Cacna1b* e37a expression might reflect an underlying alteration in the local methylation state of *Cacna1b* e37a locus. We used the spared nerve injury (SNI) model of neuropathic pain which involves ligating and transecting two of three branches of the sciatic nerve (tibial and common peroneal) on one side of the animal, while leaving the sural nerve intact (Decosterd and Woolf, 2000). The SNI model results in stable, long-term hypersensitivity in the ipsilateral area of the hind paw which is innervated by the spared sural. We measured behavioral responses in hind paw, 5-mC and *Cacna1b* e37a levels in DRG, ipsilateral and contralateral to the surgery within the same animals. Hyperalgesia in ipsilateral hind paws developed within 2 days, and was sustained at least 8 days following SNI (Fig. 7A, 7B).

**Figure 7.**
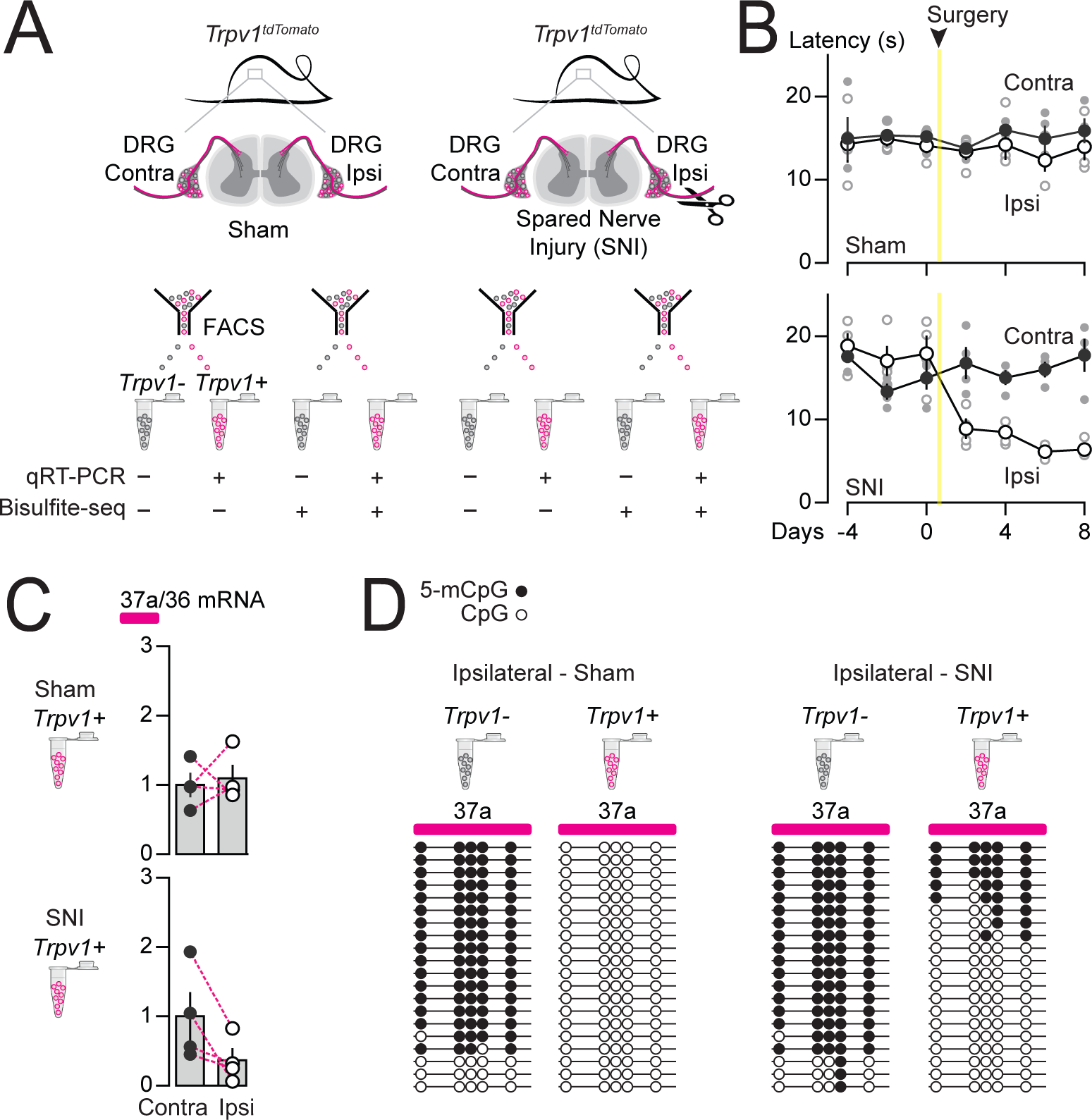
Methylation in *Cacna1b* e37a locus increased and e37a inclusion decreased in *Trpv1*-lineage neurons following peripheral nerve injury. (A) Spared nerve injury (SNI) model was used to induce prolonged hyperalgesia in *Trpv1*^*TdTomato*^ mice. Behavioral assessments are shown in *B*. *Trpv1*-lineage and non *Trpv1*-lineage cells of contralateral and ipsilateral L3 and L4 DRG from Sham and SNI mice were separated by FACS. *Trpv1*-lineage neurons were analyzed by qRT-PCR in *C*, and *Trpv1*-lineage and non *Trpv1*-lineage cells pooled from 3 mice per condition for bisulfite sequencing in *D*. (B) Thermal sensitivity assessment of mice prior to, and 8 days after surgery. Plantar, hind paw withdrawal latencies measured in response to a radiant heat source both ipsilateral and contralateral to the site of surgery in sham and SNI mice. SNI mice developed hyperalgesia in ipsilateral but not contralateral hind paws (two-way ANOVA *P* value = 0.0173 for side to the surgery factor, and *P* value = <0.0001 for time factor). By contrast, sham surgery had no effect on paw withdrawal latencies in response to heat source (two-way ANOVA *P* value = 0.2136 for side to the surgery factor, and *P* value = 0.7780 for time factor) (n = 4 mice per condition). (C) qRT-PCR of e37a relative to e36 in *Trpv1*-lineage neurons. Dotted lines connect contra and ipsilateral L3 and L4 DRG values from the same animals (Paired t-test *P* values for contra vs ipsilateral DRG = 0.3671 from sham, and *P* value = 0.0384 from SNI mice) (n = 4 mice per condition). (D) Methylation state of 5 CpG sites in *Cacna1b* e37a locus in non *Trpv1*-lineage and in *Trpv1*-lineage neurons from L3 and L4 DRG ipsilateral to the site of surgery for sham (*left*) and SNI (*right*) mice. Methylated (5-mCpG, filled circle) and unmethylated (CpG, open circle) sites are shown for independent clones ordered from most to least methylated. Each set of sequences represents data pooled from L3 and L4 DRG from 3 animals, for a total of 6 animals per condition.

We found that *Cacna1b* e37a mRNA levels were reduced in *Trpv1*-lineage neurons isolated from L3-L4 DRG ipsilateral to the injured nerve in all four animals, relative to contralateral L3-L4 DRG (Fig. 7C). By contrast, there were no consistent changes in *Cacna1b* e37a mRNA levels in *Trpv1*-lineage neurons isolated from sham DRG compared to contralateral (Fig 7C). We quantified 5-mC levels within *Cacna1b* e37a locus in *Trpv1*-lineage neurons and non *Trpv1*-lineage cells from ipsilateral L3-L5 DRG pooled from 3 sham and 3 SNI animals. We confirmed that *Cacna1b* e37a locus is hypomethylated in *Trpv1*-lineage neurons relative to the non *Trpv1*-lineage cell population in DRG (also see Fig. 6) but, after injury, 5-mC CpG at *Cacna1b* e37a locus is greater in ipsilateral *Trpv1*-lineage neurons (Fig. 7D). 5-mC CpG levels are unchanged in ipsilateral non *Trpv1*-lineage cell populations of DRG from SNI animals compared to sham (see Fig. 7D).

We conclude that 5-mC levels of CpG sites in *Cacna1b* e37a locus suppresses e37a inclusion during *Cacna1b* alternative pre-mRNA splicing *in vivo* in most neurons. Methylation of *Cacna1b* e37a locus occludes CTCF binding and accounts for the low expression of *Cacna1b* e37a mRNAs throughout the nervous system. In contrast, in *Trpv1*-lineage neurons, *Cacna1b* e37a locus is hypomethylated, permitting CTCF to bind and to promote *Cacna1b* e37a inclusion. Differential local methylation dictates the cell-specific pattern of *Cacna1b* e37a expression particularly in *Trpv1*-lineage neurons of DRG. We showed that the increase in 5-mC at CpG sites in *Cacna1b* e37a locus following peripheral nerve injury, that induces long-lasting hyperalgesia, is associated with reduced *Cacna1b* e37a expression in *Trpv1*-lineage neurons.

## DISCUSSION

Cell-specific alternative splicing is essential for normal cell function, and ~30% of all disease-causing mutations are related to RNA splicing (Montes et al., 2019). Our studies reveal the cell-specific mechanisms that govern alternative splicing of a critical synaptic calcium ion channel gene and they shed light on chronic pathology that follows nerve injury.

Cell-specific *Cacna1b* e37a splicing in *Trpv1*-lineage nociceptors underlies the unique sensitivity of voltage-gated Ca_V_2.2 calcium channels to GPCRs important for defining their properties *in vivo*. Contrary to our initial expectations, we found that a ubiquitous DNA binding protein, CTCF promotes e37a inclusion in *Cacna1b* mRNAs in a highly cell-specific pattern. This unexpected cell-specific action of CTCF is conferred by cell-specific hypomethylation within the CTCF binding motif in *Cacna1b* e37a locus. E37a locus demethylation is disrupted following nerve injury resulting in increased methylation levels and impaired *Cacna1b* e37a splicing in *Trpv1*-lineage nociceptors. Injury-induced increased methylation levels of *Cacna1b* e37a locus likely underlies some of the chronic pathophysiology associated with peripheral nerve injury.

### CTCF and methylation regulate cell-specific alternative splicing

To date, the action of CTCF on alternative splicing has only been studied in immune cells and cell lines (Agirre et al., 2015; Ruiz-Velasco et al., 2017; Shukla et al., 2011), and a role in cell-specific splicing in neurons have not been demonstrated. Our data are consistent with a model in which local hypomethylation of *Cacna1b* e37a locus permits CTCF binding, and this converts a normally weak exon, into one that is recognized by the spliceosome (Shukla et al., 2011). This splicing enhancer model for CTCF action, proposed by Oberdoerffer and colleagues studying its role in alternative splicing in *CD45* (Shukla et al., 2011), is supported by our previous analyses. For example, using two novel exon-substituted *Cacna1b* knock in mouse strains (*Cacna1b*^e37a/e37a^ and *Cacna1b*^e37b/e37b^) (Andrade et al., 2010) we observed *Cacna1b* e37a expression patterns consistent with the presence of a strong splice enhancer within *Cacna1b* e37b, except in *Tprv1*-lineage neurons where CTCF binding promotes *Cacna1b* e37a recognition and inclusion.

CTCF is proposed to enhance exon recognition by slowing the elongation rate of Pol II (Shukla et al., 2011) and this co-transcription-splicing mechanism has been proposed to account for the influence of epigenetic modifications on pre mRNA splicing in neurons (Ding et al., 2017; Schor et al., 2013). CTCF binds ~60,000 sites on average on mammalian chromosomes (Maurano et al., 2015), most of which (~70%) reside in intergenic or near to promoter regions important for regulating chromatin structure and gene expression (Barski et al., 2007; Kim et al., 2007; Ong and Corces, 2014). CTCF binding to intragenic sites is reported to be relatively invariant across different cell types (>97%) (Lee et al., 2012). But, as we document for *Cacna1b* e37a and based on analyses of cell lines, intragenic CTCF binding is variable across cell types and correlated with alternatively spliced exons (Agirre et al., 2015; Ruiz-Velasco et al., 2017; Shukla et al., 2011). Our study is consistent with the actions of CTCF controlling alternative pre mRNA splicing in immune cells and cell lines (Agirre et al., 2015; Ruiz-Velasco et al., 2017; Shukla et al., 2011), and it points to a physiological role for CTCF in cell-specific alternative splicing in neurons.

### Functional consequences of CTCF-demethylation controlled splicing

*Cacna1b* e37a is functionally important and cell-specific. Ca_V_2.2 channels containing e37a are trafficked more efficiently to the neuronal cell surface and are inhibited more effectively by G_i/o_ coupled receptors, including μ-opioid receptors (Andrade et al., 2010; Castiglioni et al., 2006; Gandini et al., 2019; Macabuag and Dolphin, 2015; Raingo et al., 2007). GPCR inhibition of Ca_V_2.2 channels at nociceptor terminals accounts for the analgesic actions of morphine at the level of the spinal cord, and cell-specific *Cacna1b* e37a inclusion translates into greater efficiency of morphine action *in vivo* (Andrade et al., 2010). After peripheral nerve injury *Cacna1b* e37a splicing in DRG is disrupted and morphine efficacy is reduced *in vivo* (Altier et al., 2007; Jiang et al., 2013). In this study we identify the underlying molecular mechanism: peripheral nerve injury leads to increased methylation levels at CpG sites within *Cacna1b* e37a locus impairing cell-specific *Cacna1b* e37a splicing.

Peripheral nerve injury triggers dynamic chromatin accessibility in DRG neurons associated with changes in gene expression and gene regions enriched with CTCF binding motifs (Palmisano et al., 2019) possibility also influencing epigenetic modifications. DNMT3a is a strong candidate for mediating increased levels of methylation of *Cacna1b* e37a locus in *Trpv1*-lineage neurons. DNMT3a is upregulated in models of neuropathic pain and is associated with hypermethylation at promotor regions of neuronal genes including *Oprm1* and *Kcna2* (Mo et al., 2018; Zhao et al., 2017); we show that DNMT3a, but not DNMT1 or DNMT3b, shifts the pattern of *Cacna1b* e37a splicing in F11 cells. Global and highly localized changes in TET1 can also modify the methylation levels of exons (Liu et al., 2016; Liu et al., 2018; Marina et al., 2016) and, as we show for dCAS9-TET1 targeted to *Cacna1b* e37a, can promote the action of CTCF to shift the pattern of alternative splicing. dCAS9-TET1 has been used to correct disease-induced alterations in gene expression and mis-splicing events (Liu et al., 2016; Liu et al., 2018), raising the possibility that this approach could be used as a strategy to correct cell-specific alternative splicing of *Cacna1b* e37a *in vivo* following peripheral nerve injury.

This study reveals a novel mechanism of cell-specific alternative splicing in neurons. Remarkably, the ubiquitous DNA binding protein, CTCF, is critical for cell-specific inclusion of a mutually exclusive exon during *Cacna1b* pre mRNA splicing in neurons. The specificity of exon inclusion arises from a striking cell-specific exon methylation pattern, providing an exciting path for understanding cell-specific CTCF action and potentially identifying a major mechanism in neuronal alternative splicing.

## Methods

See the STAR methods section.

## Supporting information

Reagent list

STAR methods

## Author contributions

EJLS & DL wrote the manuscript, designed experiments and performed analyses. EJLS performed all experiments.

## Acknowledgements

The authors thank Dr. Jennifer Pan for early discussions that guided us to CTCF as a possible regulator of alternative pre-mRNA splicing. This work was funded by grants NS055251 (D.L.) and Warren Alpert Fellowship Award (E.J.L.S).

---

**Supplementary Figure 1.**
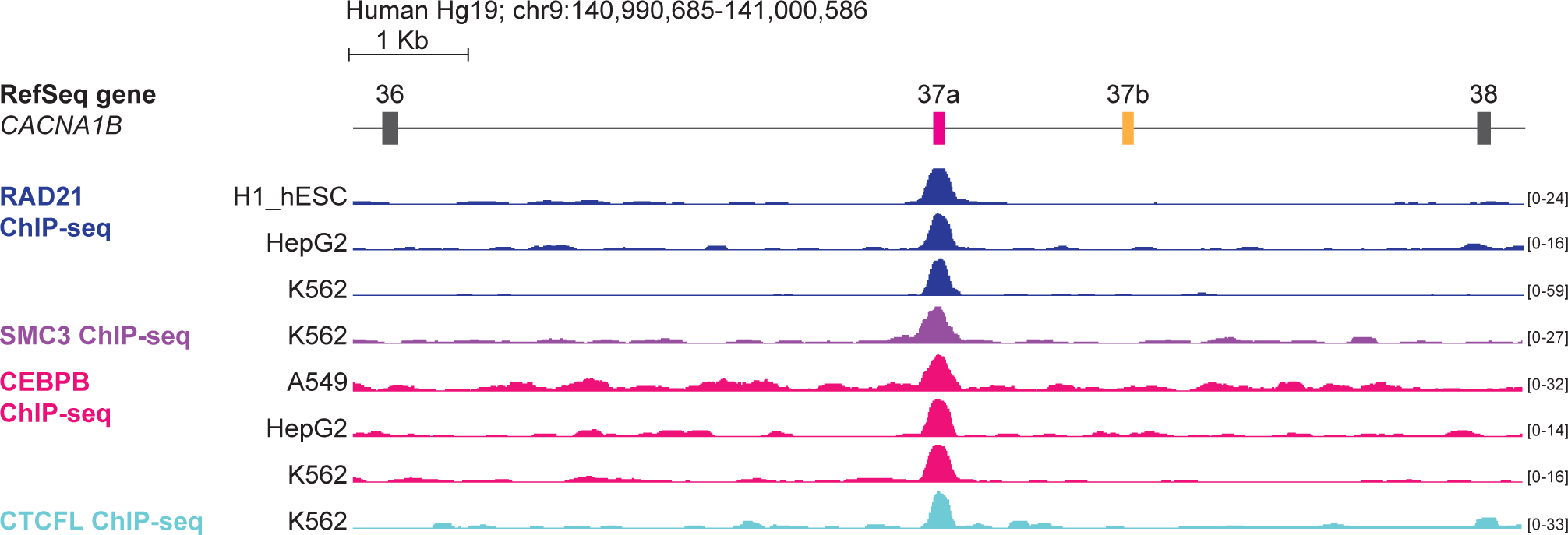
The DNA binding proteins RAD21, SMC3, CEBPB and CTCFL bind *CACNA1B* e37a locus in a small number of human cell lines. ChIP-seq signals for RAD21, SMC3, CEBPB and CTCFL binding in human cell lines aligned to ~10 kb region of *CACNA1B* (Hg19; chr9: 140,990,685-141,000,586). Y-axes for ChIP-seq tracks are scaled to the maximum signal within the selected region. Tracks with positive binding signals are shown. In total, there were binding signals in *CACNA1B* e37a locus for RAD21 in 3 of 27 cell lines, SMC3 in 1 of 27 cell lines, CEBPB in 3 of 27 cell lines, and CTCFL in 1 of 27 cell lines (https://genome.ucsc.edu/s/ejlopezsoto/Cacna1b%20e35%20to%20e38%20conservation%20track) (Consortium, 2012).

**Supplementary Figure 2.**
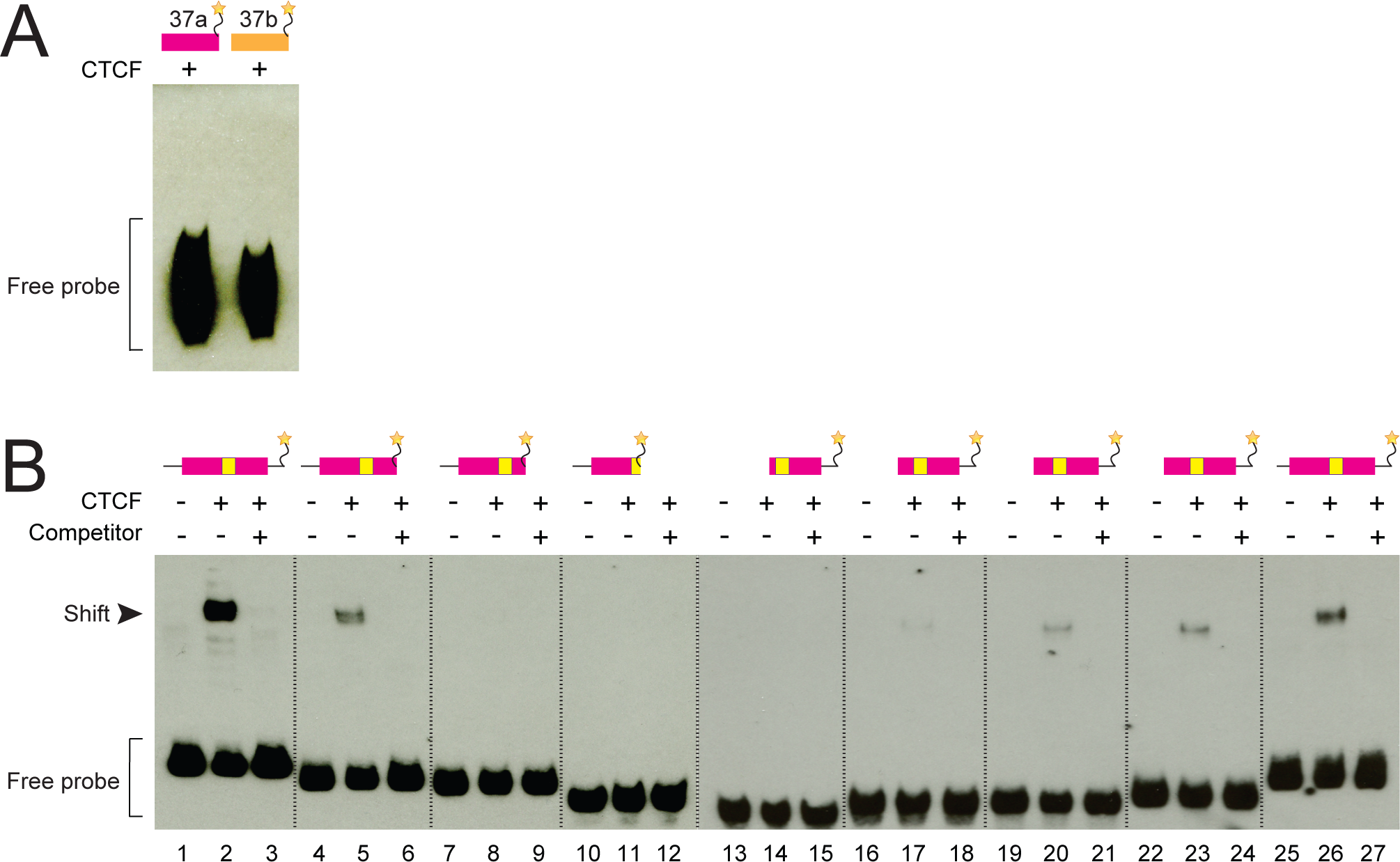
CTCF does not bind RNA *Cacna1b* e37a and CTCF binding to DNA *Cacna1b* e37a *in vitro* increases with probe length. (A) RNA gel shift assay for *Cacna1b* probes (97 bp) containing e37a or e37b with 50 ng recombinant CTCF. (B) DNA gel shift assay for different length e37a *Cacna1b* DNA probes. Full length, 3’ truncated, and 5’ truncated DNA probes containing e37a either alone (lanes 1, 4, 7, 10, 13, 16, 19, 22 and 25) or preincubated with 50 ng recombinant CTCF (lanes 2, 5, 8, 11, 14, 17, 20, 23 and 26), or CTCF plus 1000-fold excess unlabeled e37a competitor (lanes 3, 6, 9, 12, 15, 18, 21, 24 and 27). *Cacna1b* probes containing e37a (purple) and CTCF binding motif (yellow) are shown. 3’ truncated DNA probes are shown in lanes 4 to 12, and 5’ truncated DNA probes in lanes 13 to 27.

**Supplementary Figure 3.**
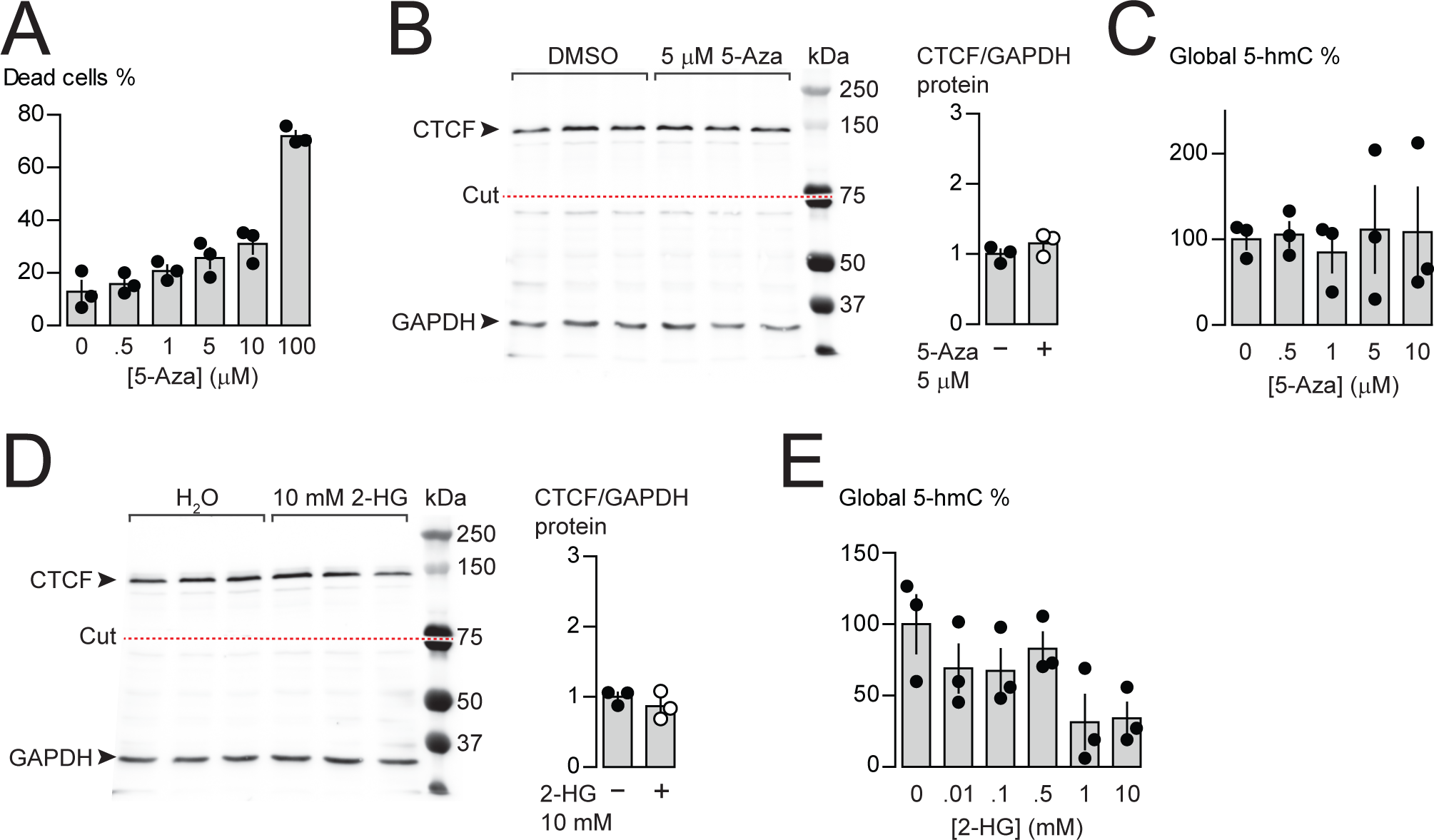
Effects of 5-Azacytidine and 2-hydroxyglutarate on cytotoxicity, CTCF and 5-hmC levels in F11 cells. (A) F11 cytotoxicity with different concentrations of 5-Azacytidine (5-Aza) for 3 days (ANOVA *P* value = <0.0001, Dunnett’s multicomparison *P* values for 0 vs 0.5 μM = 0.9380, 0 vs 1 μM = 0.3112, 0 vs 5 μM = 0.0518, 0 vs 10 μM = 0.0061, 0 vs 100 μM = <0.0001) (n= 3 per condition). (B) CTCF protein levels in F11 cells after 3 days of control (DMSO) or 5 μM 5-Aza treatment. Anti-CTCF identifies a single band at ~135 kDa. Transfer membrane was cut at ~ 75 kDa (dotted red line) and the lower part treated with anti-GAPDH to measure GAPDH levels for protein expression and loading reference. CTCF protein levels relative to GAPDH (t-test *P* value = 0.2637 for DMSO vs 5-Aza) (n = 3 per condition). (C) Global 5-hydroxymethylcytosine (5-hmC) levels in F11 cells after 3 days exposure to different concentrations of 5-Aza (ANOVA *P* value = 0.9832, Dunnett’s multicomparison *P* values for 0 vs 0.5 μM = 0.9998, 0 vs 1 μM = 0.9935, 0 vs 5 μM = 0.9975, and 0 vs 10 μM = 0.99971) (n= 3 per condition). (D) CTCF protein levels in F11 cells after 1 day of control (H_2_O) or 10 mM 2-hydroxyglutarate (2-HG) treatment. Anti-CTCF identifies a single band at ~135 kDa. Transfer membrane was cut at ~ 75 kDa (dotted red line) and the lower part treated with anti-GAPDH to measure GAPDH levels for protein expression and loading reference. CTCF protein levels relative to GAPDH (t-test *P* value = 0.3716 for H_2_O vs 2-HG) (n = 3 per condition). (E) Global 5-hmC levels in F11 cells after 1 day exposure to different concentrations of 2-HG (ANOVA *P* value = 0.0667, Dunnett’s multicomparison *P* values for 0 vs 0.01 mM = 0.5444, 0 vs 0.1 mM = 0.4971, 0 vs 0.5 mM = 0.9087, 0 vs 1 mM = 0.0417, and 0 vs 1 mM = 0.0511) (n= 3 per condition). Biological replicates represent independent cell cultures, treatment and transfections.

