## Supplementary material for "Cell-specific exon methylation and CTCF binding in neurons regulates calcium ion channel splicing and function": Reagent list

| REAGENT OR RESOURCE | SOURCE | IDENTIFIER |
| --- | --- | --- |
| **Antibodies** | | |
| CTCF (D31H2) XP® Rabbit mAb (1:1000) | Cell Signaling Technology | Cat# 3418, RRID:AB_2086791 |
| GAPDH (14C10) Rabbit mAb (1:1000) | Cell Signaling Technology | Cat# 2118, RRID:AB_561053 |
| Normal rabbit IgG | Cell Signaling Technology | Cat# 2729, RRID:AB_1031062 |
| Purified Mouse Anti-CTCF Clone 48/CTCF (RUO) | BD Biosciences | Cat# 612148, RRID:AB_399519 |
| Horseradish peroxidase-labeled anti–rabbit secondary antibody | Kirkegaard & Perry Laboratories | Cat# 474-1516 |
| **Chemicals, Peptides, and Recombinant Proteins** | | |
| CTCF human recombinant protein | Abnova | Cat# H00010664-P01 |
| Isoflurane | Patterson Veterinary | Cat# 14043070406 |
| Isopropyl alcohol | Dynarex | Cat# 1113 |
| Povidone-iodine solution | Dynarex | Cat# 1108 |
| HBSS | Gibco | Cat# 24020117 |
| Collagenase | Sigma-Aldrich | Cat# C9891 |
| Trypsin | Sigma-Aldrich | Cat# 85450C |
| PBS | Gibco | Cat# 10010031 |
| Fetal Bovine Serum | Gibco | Cat# A3160601 |
| TRIzol® LS Reagent | Invitrogen | Cat# 10296010 |
| Dulbecco’s modified Eagle’s medium | Gibco | Cat# 10569010 |
| Lipofectamine™ 2000 Transfection Reagent | Invitrogen | Cat# 11668027 |
| Opti-MEM™ reduced serum medium | Gibco | Cat# 31985070 |
| cOmplete™, Mini, EDTA-free Protease Inhibitor Cocktail | Roche | Cat# 04693159001 |
| Precision Plus Protein™ Dual Color Standards | Bio-Rad | Cat# 1610374 |
| Precision Protein™ StrepTactin-HRP Conjugate | Bio-Rad | Cat# 1610381 |
| ProSignal® Dura ECL Reagent | Genesee Scientific | Cat# 20-301 |
| EpiMark® Hot Start Taq DNA Polymerase and reaction buffer | New England Biolabs | Cat# M0490S |
| SeaKem® LE agarose | Lonza | Cat# 50004 |
| One Shot™ Stbl3™ Chemically Competent *E. coli* | Thermo Fisher Scientific | Cat# C737303 |
| Magna ChIP™ Protein A+G Magnetic Beads | Millipore | Cat# 16-663 |
| RNase Cocktail™ Enzyme Mix | Invitrogen | Cat# AM2286 |
| Proteinase K | Ambion | Cat# AM2546 |
| Q5® High-Fidelity DNA Polymerase and reaction buffer | New England Biolabs | Cat# M0491 |
| TRIzol | Invitrogen | Cat# 15596018 |

| | | |
|---|---|---|
| AarI restriction enzime | Thermo Fisher Scientific | Cat# ER1581 |
| NEBuffer™ 3.1 | New England Biolabs | Cat# B7203S |
| T4 DNA Ligase and buffer | New England Biolabs | Cat# M0202S |
| One Shot™ TOP10 Electrocomp™ *E. coli* | Thermo Fisher Scientific | Cat# C404050 |
| 5-Azacytidine | Sigma-Aldrich | Cat# A2385 |
| (2S)-2-Hydroxyglutaric Acid Octyl Ester Sodium Salt | TRC Toronto Research Chemicals | Cat# H942596 |
| **Critical Commercial Assays** | | |
| Pierce™ BCA Protein Assay Kit | Thermo Fisher Scientific | Cat# 23227 |
| QIAamp DNA Mini Kit | QIAGEN | Cat# 51304 |
| EpiTect Bisulfite Kit | QIAGEN | Cat# 59104 |
| QIAquick gel extraction kit | QIAGEN | Cat# 28704 |
| CloneJET PCR Cloning Kit | Thermo Fisher Scientific | Cat# K1232 |
| QIAprep Spin Miniprep Kit | QIAGEN | Cat# 27106 |
| QIAquick PCR Purification Kit | QIAGEN | Cat# 28106 |
| Pierce™ Biotin 3' End DNA Labeling Kit | Thermo Fisher Scientific | Cat# 89818 |
| LightShift Chemiluminescent EMSA Kit | Thermo Fisher Scientific | Cat# 20148 |
| SuperScript® III First-Strand Synthesis System with Poli-dT primers | Invitrogen | Cat# 18080051 |
| MethylFlash Methylated DNA 5-mC Quantification Kit | EpiGentek | Cat# P-1030 |
| MethylFlash Global DNA Hydroxymethylation 5-hmC ELISA Easy Kit | EpiGentek | Cat# P-1032 |
| **Experimental Models: Organisms/Strains** | | |
| B6.129-Trpv1tm1(cre)Bbm/J Mus musculus | The Jackson Laboratory | Cat# JAX:017769, RRID:IMSR_JAX:017769 |
| B6;129S-Gt(ROSA)26Sortm32(CAG-COP4*H134R/EYFP)Hze/J Mus musculus | The Jackson Laboratory | Cat# JAX:012569, RRID:IMSR_JAX:012569 |
| B6;129S6-Gt(ROSA)26Sortm14(CAG-tdTomato)Hze/J Mus musculus | The Jackson Laboratory | Cat# JAX:007908, RRID:IMSR_JAX:007908 |
| **Recombinant DNA** | | |
| pcDNA3-Tet1 | Addgene | Plasmid #60938, RRID:Addgene_60938 |
| pcDNA3-Tet2 | Addgene | Plasmid #60939, RRID:Addgene_60939 |
| pcDNA-Flag-Tet3 | Addgene | Plasmid #60940, RRID:Addgene_60940 |
| Fuw-dCas9-Tet1CD | Addgene | Plasmid #84475, RRID:Addgene_84475 |

| | | |
|---|---|---|
| pgRNA-modified | Addgene | Plasmid #84477, RRID:Addgene_84477 |
| rbCTCF-GFP | Burke et al., 2005 | N/A |
| GFP | Burke et al., 2005 | N/A |
| **Sequence-Based Reagents** | | |
| siGENOME Non-Targeting siRNA Pool #1 | GE Healthcare Dharmacon | Cat# D-001206-13 |
| SMARTpool: siGENOME Dnmt1 siRNA | GE Healthcare Dharmacon | Cat# M-056796-01 |
| SMARTpool: siGENOME Dnmt3a siRNA | GE Healthcare Dharmacon | Cat# M-065433-01 |
| SMARTpool: siGENOME Dnmt3b siRNA | GE Healthcare Dharmacon | Cat# M-044164-01 |
| SMARTpool: siGENOME Ctcf siRNA | GE Healthcare Dharmacon | Cat# M-044693-01 |
| **Primers: sequence** | **Sequence** | **Target** |
| Fw-e37a: ACCTGTAACATTTCCTTTCCAG | This paper | Fw *Cacna1b* e37a locus |
| Rv-e37a: GAGGCTCTGAAGTTGCAAAC | This paper | Rv *Cacna1b* e37a locus |
| Fw-e37b: CCTCTGGAACGGGTTTCCAG | This paper | Fw *Cacna1b* e37b locus |
| Rv-e37b: TCAGTGCAGGGTCAAGGTCTAC | This paper | Rv *Cacna1b* e37b locus |
| JLS19: TTGTTGCTGTAATCATGGACAA | This paper | Fw *Cacna1b* e36 |
| JLS20: CAGCCCAGACTCGAATGAAT | This paper | Rv *Cacna1b* e36 |
| JLS09: CGCAATACAACGCAACAAAC | This paper | Rv *Cacna1b* e37a |
| JLS10: GAGGTGGGGACATGTGTTTC | This paper | Rv *Cacna1b* e37b |
| JLS21: AATGTGTCCGTCGTGGATCT | Toyoda et al., 2014 | Fw *Gadph* |
| JSL22: GTTGAAGTCGCAGGAGACAA | Toyoda et al., 2014 | Rv *Gadph* |
| JLS47: TATTTTTTATTGTAGATTGGGTGGG | This paper | Fw bisulfite converted *Cacna1b* e37a locus |
| JLS48: TCAAATAAAAACTCTAAAATTACAAA | This paper | Rv bisulfite converted *Cacna1b* e37a locus |
| JLS53: CGACTCACTATAGGGAGAGCGGC | Universal primer | Fw pJET1.2; for sequencing bisulfite converted *Cacna1b* e37a |
| JLS54: AAGAACATCGATTTTCCATGGCAG | Universal primer | Rv pJET1.2; for sequencing bisulfite converted *Cacna1b* e37a |
| JLS65: GAAACTCACCCTAACTG | This paper | For sequencing pgRNA plasmid insert |
| JLS59: TTGGACCTTGTAGGCCAACCTACG | This paper | Fw oligo for gRNA1 e37a |
| JLS60: AAACCGTAGGTTGGCCTACAAGGT | This paper | Rv oligo for gRNA1 e37a |
| JLS61: TTGGCAGTTGCCGGATTCATTATA | This paper | Fw oligo for gRNA2 e37a |
| JLS62: AAACTATAATGAATCCGGCAACTG | This paper | Rv oligo for gRNA2 e37a |

| | | |
|---|---|---|
| JLS63: TTGGCCCCGGGGAAAAATTTTTTT | This paper | Fw oligo for gRNAnt e37a |
| JLS64: AAACAAAAAAATTTTTCCCCGGGG | This paper | Rv oligo for gRNAnt e37a |
| **Software and Algorithms** | | |
| MethPrimer | Li and Dahiya, 2002 | URL: http://urogene.org/; RRID:SCR_010269 |
| Primer3 | Koressaar and Remm, 2007 | URL: http://primer3.ut.ee; RRID:SCR_003139 |
| ImageJ | NIH | URL: https://imagej.net/; RRID:SCR_003070 |
| Prism 8 | GraphPad | URL: http://www.graphpad.com/scientific-software/prism/; RRID: SCR_005375 |
| **Other** | | |
| Sterile 6-0 coated vicryl suture | Ethicon | Cat# J833G |
| Deltaphase® Isothermal Pads | Braintree scientific | Cat# DPIP |
| Plantar Analgesia Meter | IITC | Cat# II-390G |
| Flowmi® Cell Strainer | Sigma-Aldrich | Cat# BAH136800070 |
| Amersham Protan 0,45 mM NC, nitrocellulose membrane | GH Healthcare Life Science | Cat# 10600002 |
| Amersham™ Hybond™-N+ 0.45 µm nitrocellulose membrane | GH Healthcare Life Science | Cat# 95038-376 |
