## Supplementary material for "Cell-specific exon methylation and CTCF binding in neurons regulates calcium ion channel splicing and function": STAR methods

### CONTACT FOR REAGENT AND RESOURCE SHARING

Information and requests for resources and reagents should be directed to and will be fulfilled by the Lead Contact Diane Lipscombe. There are no restrictions on any data or materials presented in this paper.

### EXPERIMENTAL MODEL AND SUBJECT DETAILS

#### Mice

Mice were housed and bred at Brown University. All protocols and procedures were approved by the Brown University Institutional Animal Care and Use Committee. Mice were maintained at 22°C in a 12 h light/dark cycle, with food and water available *ad libitum*. *Trpv1*$^{Cre}$ (Cavanaugh et al., 2011) (Cat# JAX:017769, RRID:IMSR_JAX:017769), *lox-STOP-lox*$^{ChR2\text{-}EYFP}$ (Cat# JAX:012569, RRID:IMSR_JAX:012569) and *lox-STOP-lox*$^{TdTomato}$ (Madisen et al., 2012) (Cat# JAX:007908, RRID:IMSR_JAX:007908) mice were purchased from The Jackson Laboratory. Mice used in this study were first generation double heterozygous offspring from single homozygous parents. All procedures used to generate data reported in this study are described here. Mice had no prior history of drug administration, surgery or behavioral testing. Behavioral experiments were performed on male and female mice 3-5 month of age.

### METHOD DETAILS

#### Spared Nerve injury

The spared nerve injury (SNI) mouse model (Decosterd and Woolf, 2000; Jiang et al., 2013) was performed on male and female mice 3-5 month of age to develop long lasting neuropathic pain. Mice were anesthetized with 3% isoflurane (induction, Patterson Veterinary Cat# 14043070406) and 2% isoflurane (maintenance) with oxygenation throughout surgery. The left hind leg was shaved, wiped clean with sterile 70% isopropyl alcohol (Dynarex Cat# 1113) and sterile 10% povidone-iodine solution (Dynarex Cat# 1108). An ~1 cm incision was made in the skin of the upper thigh, approximately where the sciatic nerve trifurcates. The common peroneal, tibial, and sural branches of the sciatic nerve were exposed by blunt dissection and teasing open biceps femoris muscle. The common peroneal and tibial nerves were ligated with a sterile 6-0 coated vicryl suture (Ethicon Cat# J833G) and transected 2 mm from the distal end. The surgical field was irrigated with sterile saline, the skin incision sutured closed, and then wiped clean with sterile 10% povidone-iodine solution (Dynarex Cat# 1108). For sham-operated mice, the surgery was identical except that the nerves were not ligated or transected after exposure. Immediately after surgery, mice were placed on a warm Deltaphase® Isothermal Pads (Braintree scientific Cat# DPIP) to recover for ~1hr before being returned to their home cage. Mice were closely monitored for three days.

**Thermosensitivity assessment**

We used a Plantar Analgesia Meter (IITC Cat# II-390G) to assess thermal responses to radiant heat from a focal heat source (Hargreaves method) (Hargreaves et al., 1988). All testing was performed at room temperature (22°C). Mice were habituated for at least 30 min in 4 individual Plexiglas chambers placed on a 22 cm elevated glass stage. Males and females were tested in separate trials. A focused visible-light, radiant heat source was positioned beneath the plantar surface of the hind paw. In all experiments, the high-intensity beam was set at 40%, low-intensity beam set at 10%, and maximum trial duration at 30s. An orange-pass filter was used to prevent blue-light activation of mice expressing channelrhodopsin in sensory nerve terminals. Paw withdrawal latencies were measured in response to radiant heat source applied to the plantar surface of the hind paw. We assessed withdrawal latencies for contralateral and ipsilateral paws, before and after surgery, in SNI and sham mice. We measured latencies in response to three individual trials for each hind paw, and trials were separated by at least 2 mins. Withdrawal responses were counted if the hind paw withdrawal was rapid and associated with licking or shaking. Withdrawal thresholds were calculated as the average withdrawal latency for the three trials. The experimenter was blind to genotype and injury, and each testing group (4 mice) was randomized for SNI and sham conditions.

**DRG dissociation and cell sorting**

Freshly dissociated DRG were prepared from 3-5 month male and female mice. Under isoflurane anesthesia, mice were euthanized and all axial DRG (figure 6) or L3-L4 DRG (figure 7) removed. Total dissection time was kept under 30 min. Freshly dissected DRG were placed into ice-cold HBSS (Gibco Cat# 24020117), roots were carefully removed and briefly washed in HBSS (Gibco Cat# 24020117). DRG were enzymatically treated with 2 mg/mL collagenase (Sigma-Aldrich Cat# C9891) and 0.25% trypsin (Sigma-Aldrich Cat# 85450C) in HBSS (Gibco Cat# 24020117) at 37°C for 20 mins, and separated by gently mechanical dissociation using a pipette tip. Cells were centrifuged at 300 g for 7 minutes at RT°C, resuspended in PBS (Gibco Cat# 10010031) and passed through a Flowmi® Cell Strainer (porosity 70 μm, Sigma-Aldrich Cat# BAH136800070) pre-rinsed with PBS. Cells were collected and centrifuged at 300 g for 7 minutes at 4°C, resuspended in 1-3 mL PBS (Gibco Cat# 10010031) with 10% FBS (Gibco Cat# A3160601) and kept on ice. Samples were immediately transported to the Flow Cytometry and Sorting Facility at Brown University. DRG cells were separated into *Trpv1*-lineage and non-*Trpv1*-lineage by fluorescent-activated cell sorting using a BD FACS Aria IIIu cytometer and sorter. Sorting criteria were determine monitoring distribution of GFP or TDTOMATO signals; a subsample of sorted cells were examined under fluorescent microscope for validation of sorting. At least 5000 neurons were collected for each sample in PBS (Gibco Cat# 10010031) with 10% FBS (Gibco Cat# A3160601) when used for gDNA extraction or TRIzol® LS Reagent (Invitrogen Cat# 10296010) when used for RNA extraction. Samples were sorted at 4°C and kept on ice until used within 1 hr.

**Cell culture**

The F11 cell line is a somatic cell hybrid of rat embryonic DRG and mouse neuroblastoma cell line N18TG2. F11 cells express receptors and ion channels found in nociceptors including Cav2.2 channels (Allen et al., 2017; Yin et al., 2016). F11 cell cultures were maintained at sub-confluent density in Dulbecco’s modified Eagle’s medium (Gibco Cat# 10569010) with 10% FBS (Gibco Cat# A3160601). Exponentially dividing cultures were transfected using Lipofectamine™ 2000 Transfection Reagent (Invitrogen Cat# 11668027) and Opti-MEM™ reduced serum medium (Gibco Cat# 31985070) as per manufacturer’s protocol.

**Western blot**

Dissociated F11 cells were lysed in ice-cold RIPA buffer (25mM Tris•HCl pH 7.6, 150mM NaCl, 1% NP-40, 1% sodium deoxycholate, 0.1% Sodium dodecyl sulfate and 1X protease inhibitor (cOmplete™, Mini, EDTA-free Protease Inhibitor Cocktail, Roche Cat# 04693159001) and kept on ice during the following procedure: 30 mins incubation, sonication (3 cycles @ 30 sec on and 1 min off, at high amplitude with a Q500A sonicator system Qsonica), and 10 min centrifugation at 14000g. Proteins were detected, and concentrations determined from supernatants by Pierce™ BCA Protein Assay Kit (Thermo Fisher Scientific, cat#23227) prior to Western blotting.

For Western blotting and analysis: Protein samples were denatured in Laemmli buffer at 95°C for 5 minutes and immediately placed on ice until needed. 10 µg of each sample and 8 µl of Precision Plus Protein™ Dual Color Standards (Bio-Rad, cat#1610374) were separated in SDS-PAGE (4% stacking and 6% resolving gels). Samples were run for 1 hr at 10 mA followed by 13 min at 20 mA, and transferred to nitrocellulose membrane for 1 hr at 100 V (GH Healthcare Life Science, Amersham Protan 0,45 µM NC Cat# 10600002). Membranes were rinsed with PBS-T, blocked with 2.5% nonfat milk and 5% BSA in PBS-T for either 2 hrs at RT°C, or overnight at 4°C. Membranes were cut at ~ 75 kDa and >75 kDa region incubated in CTCF primary antibody (CTCF (D31H2) XP® Rabbit mAb - Cell Signaling Technology, Cat# 3418, RRID:AB_2086791) and <75 kDa region with GAPDH primary antibody (GAPDH (14C10) Rabbit mAb - Cell Signaling Technology, Cat# 2118, RRID:AB_561053). Primary antibodies were diluted in 5% BSA PBS-T and membranes were incubated for 2 hrs at RT°C or overnight at 4°C (CTCF 1:1000; GAPDH 1:1000). Membranes were rinse 2x in PBST-T and 4x in PBS-T for 10 min, both at RT°C, and incubated at 1 hr at RT°C with horseradish peroxidase-labeled anti–rabbit secondary antibody (1:15,000; Kirkegaard & Perry Laboratories Cat# 474-1516), and Precision Protein™ StrepTactin-HRP Conjugate (1:10,000 - Bio-Rad, cat#1610381) for ladder detection. Membranes were rinsed 2x in PBS-T and washed 3x in PBS-T for 10 min at RT°C.

Protein bands were visualized using ProSignal® Dura ECL Reagent (Genesee Scientific, cat#20-301) and imaged with an Azure C600 system (Azure Biosystems). A series of exposures were sampled to ensure that signals were in the linear range for detection. Intensities of protein bands were quantified in non-saturated images using ImageJ software (Schneider et al., 2012). CTCF protein expression levels were normalized to GAPDH levels measured from the same gel. Complete western blots are shown for each figure.

**Bisulfite sequencing**

Genomic DNA (gDNA) from F11r DRG was extracted using QIAamp DNA Mini Kit (QIAGEN Cat# 51304). Bisulfite conversion of gDNA was performed using the EpiTect Bisulfite Kit (QIAGEN Cat# 59104) following the low-concentration sample protocol according to the manufacturer's instructions. *Cacna1b* e37a bisulfite converted gDNA fragments (207 bp) were amplified by PCR in 10 µl reactions containing 4 µl of bisulfite converted gDNA; 0.3 µM forward and reverse primers (JLS47 and JLS48, table 1) designed with MethPrimer (Li and Dahiya, 2002); 200 µM dNTPs; 1.25 units EpiMark® Hot Start Taq DNA Polymerase and 1X reaction buffer (New England Biolabs Cat# M0490S); and $H_2O$ in a C1000 Touch Thermal Cycler (BIO RAD). PCR reactions consisted of 1 min of initial incubation at 95°C and 40 cycles of 30 s at 95°C, 1 min at 58°C, and 1 min at 68°C, followed by 5 min at 68°C. PCR products were visualized in a 1.5% SeaKem® LE agarose (Lonza Cat# 50004), extracted using QIAquick gel extraction kit (QIAGEN Cat# 28704) and subcloned into pJET1.2/blunt vector using CloneJET PCR Cloning Kit (Thermo Fisher Scientific Cat# K1232). One Shot™ Stbl3™ Chemically Competent *E. coli* (Thermo Fisher Scientific Cat# C737303) were transform with the ligation mixture; and at least 20 independent clones per condition were amplified overnight, isolated using QIAprep Spin Miniprep Kit (QIAGEN Cat# 27106) and sequenced using Fw or Rv pJET1.2 primers (JLS53 and JLS54, table 1). Methylation status of *Cacna1b*-e37a locus was determined by comparing bisulfite converted sequences with the original *Cacna1b*-e37a sequence. The overall efficiency of bisulfite conversion was superior to 99 % as estimated from cytosines from non-CpG sites.

**Chromatin Immunoprecipitation (ChIP)**

Protein-DNA crosslinking was performed at RT°C in F11 cells with 1% formaldehyde in PBS (Gibco cat# 10010031) for 10 min and gentle shaking. The crosslinking reaction was quenched by adding glycine (final concentration of 125 mM) during 5 min rotation mixing at RT°C.

Samples were washed 3x in chilled PBS and resuspended in lysis buffer (50 mM HEPES-KOH pH 7.5, 140 mM NaCl, 1 mM EDTA, 1% Triton X-100, 0.1% sodium deoxycholate, 0.1% sodium dodecyl sulfate and 1X protease inhibitor (cOmplete™, Mini, EDTA-free Protease Inhibitor Cocktail, Roche Cat# 04693159001)). Cells were resuspended 20x using pipette and vortex 3x during 10 sec. Chromatin sonication was performed in a Q500A sonicator system (Qsonica) using 2x 5 cycles (30 s on and 1 min off, high amplitude at 4ºC) separated by a 10 min incubation on ice. Samples were centrifuged at 4ºC for 8 min at 10,000 g. Samples were resuspended in dilution buffer (16.7 mM Tris-HCl pH 8, 1.2 mM EDTA, 334 mM NaCl, 2.2% Triton X-100, 0.01% SDS, and 1X protease inhibitor (cOmplete™, Mini, EDTA-free Protease Inhibitor Cocktail, Roche Cat# 04693159001)) to a final volume of 500 µl. 5% of the sample was removed to use as a positive control (input). Chromatin was inmunoprecipitated by adding 2 µg CTCF (D31H2) XP® Rabbit mAb (CTCF (D31H2) XP® Rabbit mAb - Cell Signaling Technology Cat# 3418, RRID:AB_2086791) or 2 µg of normal rabbit IgG (Cell Signaling Technology Cat# 2729, RRID:AB_1031062) followed by overnight incubation at 4ºC with gentle shaking. 30 µl Magna ChIP™ Protein A+G Magnetic Beads (Millipore, cat#16-663) were added to each sample and incubated 2 hs at 4ºC with gentle shaking. Prior to use, magnetic beads were washed twice with dilution buffer at RTºC to reduce background. The Ab-CTCF-DNA complex was displaced from the magnetic beads and collected after sequential washes in the following buffers for 4 min each at 4ºC: low salt buffer (20 mM Tris-HCl pH 8, 150 mM NaCl, 2 mM EDTA, 0.1% SDS, 1% Triton X-100); high salt buffer (20 mM Tris-HCl pH 8, 500 mM NaCl, 2 mM EDTA, 0.1% SDS, 1% Triton X-100); LiCl buffer (10 mM Tris-HCl pH 8, 1 mM EDTA, 250 mM LiCl, 1% NP40, 1% sodium deoxycholate); and finally TE. The IP and input samples were diluted in elution buffer (50mM Tris-HCl pH 8, 10 mM EDTA, 50 mM $NaHCO_3$ and 1% sodium dodecyl sulfate), incubated with RNase Cocktail™ Enzyme Mix (1 µg final concentration; Invitrogen Cat# AM2286) 20 min at 37ºC and further incubated with proteinase K (20 µg/ml; Ambion/Thermo Fisher Scientific Cat# AM2546) for 3 hrs at 65°C. gDNA was isolated using QIAquick PCR Purification Kit (QIAGEN Cat# 28106).

IP and input gDNAs were amplified in the real-time PCR reaction using *Cacna1b* e37a or e37b specific primer pairs (Fw-e37a and Rv-e37a for e37a; Fw-e37b and Rv-e37b for e37b, table 1) in a 10 µl reaction with 0.3 µM forward and reverse primers, 1X Power SYBR® Green PCR Master Mix (Thermo Fisher Scientific Cat # 4368706) and $H_2O$ in a StepOnePlus™ Real-Time

PCR System (Applied Biosystems). PCR reactions consisted of 10 min of initial incubation at 95°C and 45 cycles of 20 s at 95°C, and 1 min at 60°C.

Quantification of CTCF bound to *Cacna1b* e37 loci was calculated using the formula $100*2^{\Delta Ct}$ $^{(adjusted\ input\text{-}IP)}$; normalized to IgG condition.

**Electrophoretic Mobility Shift Assay (EMSA)**

DNA double-stranded *Cacna1b* e37a and e37b probes were PCR-amplified from mouse gDNA using PAGE purified specific primer pairs (Integrated DNA Technologies). gDNA fragments were amplified by PCR in 25 µl reactions containing 0.5 µg mouse gDNA; 0.5 µM forward and reverse primers (Fw-e37a and Rv-e37a for *Cacna1b* e37a locus; and Fw-e37b and Rv-e37b for *Cacna1b* e37b locus; table 1) designed with Primer3 (Koressaar and Remm, 2007); 200 µM dNTPs; 1 unit Q5® High-Fidelity DNA Polymerase and 1X reaction buffer (New England Biolabs Cat# M0491); and $H_2O$ in a C1000 Touch Thermal Cycler (BIO RAD). PCR reactions consisted of 30 s of initial incubation at 98°C and 35 cycles of 10 s at 98°C, 20 s at 65°C, and 10 s at 72°C, followed by 2 min at 72°C. PCR products were visualized in a 1% agarose gel (Lonza Cat# 50004), extracted using QIAquick gel extraction kit (QIAGEN Cat# 28704) and followed by standard ethanol precipitation. DNA probes were label using the Pierce™ Biotin 3' End DNA Labeling Kit (Thermo Fisher Scientific Cat# 89818) and biotinylation efficiency was evaluated by Dot Blot. Annealing of DNA probes was perform by following 6 min at 95 °C and a passive slow ramp until 22°C for 1 h. RNA *Cacna1b* e37a and e37b probes (90 nucleotides) were 3' biotinylated and HPLC purificated (Integrated DNA Technologies).

To determine nucleic acid-protein interaction, 20 fmol probe was incubated in binding reaction buffer (10 mM Tris, 50 mM KCl, 5 mM MgCl2, 0.1 mM ZnSO4, 1 mM DTT, 0.1% (v/v) NP-40, 50 ng/µl Poly (dl·dC) and 2.5% (v/v) glycerol) for 30 min at 25 °C with 0.1 µg CTCF human recombinant protein (Abnova Cat# H00010664-P01). To confirm specificity of binding, 0.5 µg mouse anti-CTCF (Purified Mouse Anti-CTCF Clone 48/CTCF (RUO), BD Biosciences Cat# 612148, RRID:AB_399519) or 1000-fold excess of an unlabeled DNA probe was added to the binding reaction buffer. DNA-protein complexes were separated in 5% native polyacrylamide gel using 0.5 X Tris/borate/EDTA buffer at 4°C and transferred to positively charged nitrocellulose membrane (Hybond™-N+ 0.45 μm; Amersham™ Cat# 95038-376). EMSAs were perform using the LightShift Chemiluminescent EMSA Kit (Thermo Fisher Scientific Cat# 20148).

DNA and RNA bands were imaged with an Azure C600 system (Azure Biosystems). A series of exposures were sampled to ensure that signals were in the linear range for detection. Intensities of bands were quantified in non-saturated images using ImageJ software (Schneider et al., 2012). CTCF bound to DNA probes was calculated as % of shifted band intensity normalized to total band intensity (shifted + free bands) per condition from the same blot.

**RNA extraction, RT-PCR and qPCR**

RNA extraction from F11 cells was performed using TRIzol (Invitrogen Cat# 15596018) and from dissociated DRG cells using TRIzol® LS Reagent (Invitrogen Cat# 10296010) according to the manufacturer's instructions. 1-2 μg of RNA was immediately reverse transcribed to cDNA using the SuperScript® III First-Strand Synthesis System with Poli-dT primers (Invitrogen Cat# 18080051). *Cacna1b* e36, e37a, e37b or *Gapdh* cDNAs were amplified in 10 μl real-time PCR reactions using 0.3 μM forward and reverse primers (JLS19 and JLS20 for *Cacna1b* e36; JLS19 and JLS09 for e37a; JLS19 and JLS10 for e37a; and JLS21 and JLS22 for *Gapdh* (Toyoda et al., 2014), table 1), 1X Power SYBR® Green PCR Master Mix (Thermo Fisher Scientific Cat # 4368706) and $H_2O$ in a StepOnePlus™ Real-Time PCR System (Applied Biosystems). PCR reactions consisted of 10 min of initial incubation at 95°C and 45 cycles of 20 s at 95°C, and 1 min at 60°C. Each sample was run in triplicate per target. PCR efficiencies were calculated for all primer pairs by qPCR analysis of serial dilutions of gDNA containing target sequences, and using 5-9 replicates per DNA concentration assessed. Efficiency was calculated obtaining standard curves as described in (Pfaffl, 2001). PCR specificity was determined by PCR product length using 2 % agarose electrophoresis gel, and by post PCR melting curve analysis as follows: 15 s at 95°C followed by a temperature ramp from 60°C to 95°C in 0.3°C steps every 15 s. *Cacna1b* e37 quantification was calculated by the following ratio = $(E\ e37)^{\Delta Ct\ e37\ (control\text{-}sample)} / (E\ e36)^{\Delta Ct\ e36\ (control\text{-}sample)}$; normalized to control condition.

**Genomic DNA methylation and hydroxyl-methylation ELISAs**

gDNA was extracted from F11 cells using QIAamp DNA Mini Kit (QIAGEN Cat# 51304) and global DNA methylation (5-mC) and hydroxyl-methylation (5-hmC) levels were measured by ELISA assays (MethylFlash Methylated DNA 5-mC Quantification Kit - EpiGentek Cat# P-1030; MethylFlash Global DNA Hydroxymethylation 5-hmC ELISA Easy Kit - EpiGentek Cat# P-1032) according to manufacturer's protocols.

**cDNA plasmids and siRNAs**

cDNA plasmid expression vectors are listed in table 1. pcDNA3-Tet1 (Addgene plasmid # 60938; http://n2t.net/addgene:60938; RRID:Addgene_60938), pcDNA3-Tet2 (Addgene plasmid # 60939; http://n2t.net/addgene:60939; RRID:Addgene_60939), and pcDNA-Flag-Tet3 were gifts from Yi Zhang (Addgene plasmid # 60940; http://n2t.net/addgene:60940; RRID:Addgene_60940) (Wang and Zhang, 2014). Fuw-dCas9-Tet1CD (Addgene plasmid # 84475; http://n2t.net/addgene:84475; RRID:Addgene_84475) and pgRNA-modified were gifts from Rudolf Jaenisch (Addgene plasmid # 84477; http://n2t.net/addgene:84477; RRID:Addgene_84477) (Liu et al., 2016). CTCF-GFP and GFP were gifts from Rainer Renkawitz (Burke et al., 2005).

Guide RNA (gRNA) expression constructs were cloned by inserting annealed oligo pairs JLS59-JLS60 (gRNA1); JLS61-JLS62 (gRNA2); and JLS63-JLS64 (gRNAnt) (table 1) into modified pgRNA plasmid (Addgene Cat# 84477) (Liu et al., 2016) in AarI site (Thermo Fisher Scientific Cat# ER1581). Oligo pairs were anneal in 1X NEBuffer™ 3.1 (New England Biolabs Cat# B7203S) following 4 min at 95°C, 10 min at 70°C and a passive slow ramp until 22°C. Vector and insert were ligated using T4 DNA Ligase and 1X buffer (New England Biolabs Cat# M0202S) at room temperature for 2 hours. One Shot™ TOP10 Electrocomp™ *E. coli* (Thermo Fisher Scientific Cat# C404050) were transformed with the ligation mixture; and independent clones per condition were amplified overnight, isolated using QIAprep Spin Miniprep Kit (QIAGEN Cat# 27106) and sequenced using JLS65 primer (table 1).

siRNAs are listed in table 1. *Ctcf*, *Dnmt1*, *Dnmt3a*, *Dnmt3b* and non-targeting SMARTpool siGENOME siRNAs (GE Healthcare Dharmacon Cat# M-044693-0, M-056796-01, M-065433-

01, M-044164-01, and D-001206-13) were resuspended in RNase free $H_2O$ and shacked 30 min at RT, and storage at -80°C until need them.

**Drugs**

F11 cells were treated with 5-Azacytidine (Sigma-Aldrich Cat# A2385) to inhibit DNA methyltransferase action and promote a decrease in global methylation; and (2S)-2-Hydroxyglutaric Acid Octyl Ester Sodium Salt (TRC Toronto Research Chemicals Cat# H942596) to inhibit ten eleven translocase enzymes (TET) and promote an increase in global methylation. 5-Azacytidine was dissolved in DMSO and kept on ice until use or storage at -80°C. (2S)-2-Hydroxyglutaric Acid Octyl Ester Sodium Salt was dissolved in $H_2O$ and kept on ice until use or storage at -4°C. Concentration and incubation times are indicated in each experiment.

**Statistical Analysis**

All behavioral testing was randomized and blinded. N values represent biological replicates (number of animals, individual transfections). We plot all individual biological replicates in figures and corresponding mean and SE. Experimental groups were compared using paired or unpaired t-test, two-way or one way ANOVA followed by Dunnett's test as indicated. Absolute *P* values are reported in figure legends.
